# Plasmid biology is compressed in host chromosomal architecture

**DOI:** 10.64898/2026.08.12.744350

**Authors:** Yahui Hou, Wenzhi Xue, Teng Wang

## Abstract

Plasmids drive horizontal gene transfer and the spread of antibiotic resistance, yet their distribution across microbial genomes is highly uneven and often viewed as environmentally driven, leaving unresolved whether host chromosomal architecture imposes predictable constraints. Here, using machine learning on 52,393 complete prokaryotic genomes, we show that chromosomal gene content encodes predictive information for multiple dimensions of plasmid biology: carriage status, quantitative load, and mobility potential. Remarkably, highly compressed chromosomal signatures— as few as 15 genes or the coarse-grained composition of seven major enzyme classes—suffice for robust prediction. Moreover, different functional cargoes carried by plasmids, including antibiotic resistance classes, can also be predicted from host chromosomal signatures. These findings establish that plasmid–host compatibility is systematically encoded in host chromosomes, reframing plasmid ecology from environment-driven to host-constrained—a shift with direct implications for combating resistance and engineering stable microbial chassis.

## Introduction

The evolutionary success of prokaryotes is driven not only by mutations, but also by the continual exchange of genetic information through horizontal gene transfer (HGT)^1^. Among the vehicles of HGT, plasmids play a central role by disseminating antibiotic resistance, metabolic capabilities and virulence determinants across microbial communities^2,3^. Their abundance, however, is strikingly uneven across prokaryotic genomes: some cells harbor multiple plasmids, whereas others are entirely plasmid-free^4,5^. Explaining this striking heterogeneity remains a fundamental challenge in microbial ecology and evolution, because it ultimately governs how gene flow is organized within microbial populations.

Current theories largely attribute plasmid distribution to extrinsic ecological forces^6–9^. Environmental selection^10^, including antibiotic exposure, heavy metals, nutrient availability and biotic interactions, is thought to determine whether plasmids are maintained or lost within microbial hosts (Fig. 1a). This framework has successfully explained many context-dependent dynamics of plasmid persistence^11^. However, it implicitly assumes that microbial genomes are largely passive recipients of mobile DNAs, with plasmid fate determined mainly by ecological opportunities.

**Figure 1.**
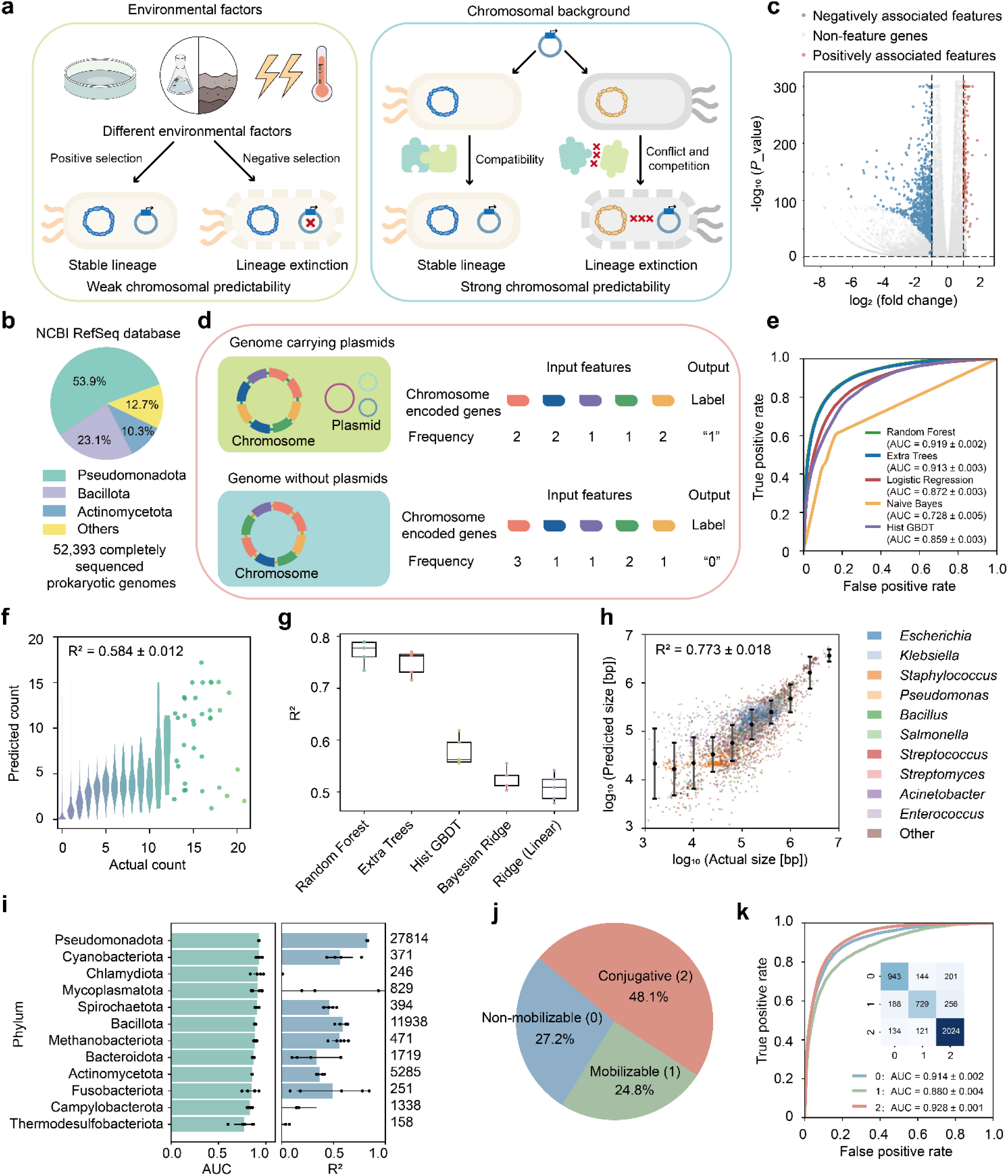
Host chromosomal architecture predicts plasmid carriage, quantitative load and mobility. (a) Conceptual framework contrasting two alternative models of plasmid ecology. Under an environment-dominated model, plasmid fate is determined mainly by ecological opportunities and selection. Thus, chromosomal gene content is expected to exhibit limited predictive power for plasmid carriage and burden (left). Under a host-genome-dominated model, plasmid presence and abundance should be systematically encoded within chromosomal functional composition and therefore be highly predictable from host chromosomal features (right). (b) Taxonomic distribution of complete prokaryotic genomes retrieved from RefSeq. (c) Identification of chromosomal feature genes based on enrichment analysis. Positively associated features are highlighted in red, negatively associated features in blue, and non-feature genes in grey. (d) Machine-learning framework using chromosomal gene-abundance profiles to predict plasmid-carriage status. (e) Comparison of classification algorithms for plasmid carriage prediction. Performance was evaluated across five independent random train–test splits. Random Forest achieved the highest predictive performance. (f) Prediction of plasmid number from chromosomal gene content. Observed and predicted plasmid counts are shown for genomes in a representative test set. (g) Comparison of regression algorithms for predicting total plasmid size. Random Forest consistently outperformed alternative models. Boxplots show the median (center line), interquartile range (box limits) and min–max values (whiskers) of scores across five train–test splits. (h) Random Forest prediction of total plasmid size using chromosomal gene content. Observed versus predicted plasmid sizes (log₁₀ bp) are shown for genomes in a representative test partition (n = 10,479). Points represent individual genomes and are colored by the ten most abundant genera (all others grouped as “Other”). Filled circles indicate mean predicted values within bins of observed plasmid size, and error bars denote ± s.d. of predicted values within each bin. The reported R² value corresponds to the mean ± s.d. across five independent train–test splits. (i) Random Forest performance across major phyla. Per-phylum classification (AUC; left) and total plasmid size regression (R²; right) performance were evaluated on test-set genomes from five independent train-test splits. Only phyla with a cumulative total of at least 100 test genomes across the five independent test splits are shown. Numbers on the right indicate the cumulative number of test genomes across the five test splits for each phylum. Bars and error bars denote the mean ± s.d. of metric scores across five replicates, with individual replicate values overlaid as dots. (j) Distribution of genomes assigned to non-mobilizable (0), mobilizable (1), and conjugative (2) classes (n = 23,717). (k) Prediction of genome-level plasmid mobility classes. One-versus-rest ROC curves showing classifier performance. Curves represent the mean true-positive rate across five independent train–test splits. The inset confusion matrix shows the element-wise mean confusion matrix across the five independent train–test splits, with values truncated to integers.

An alternative possibility is that chromosomes themselves encode the capacity to accommodate plasmids^12–14^. Plasmids often impose demands on host metabolism, gene regulation, DNA replication and protein synthesis^15,16^, suggesting that successful maintenance requires compatibility with pre-existing cellular machineries^17–19^. Consistent with this notion, the stability, conjugation efficiency and gene expression level of the same plasmid often vary among different hosts^15,20,21^. These observations imply that the chromosome may not simply provide the background in which plasmids evolve, but instead define the evolutionary landscape on which plasmid acquisition and persistence occur^22–24^. If so, the intrinsic organization of host chromosomes should contain predictive information about their ability to acquire and maintain mobile genetic elements. However, whether such predictive constraints exist, and how extensively they shape plasmid biology across prokaryotes, remains unknown.

Here we address this question by systematically interrogating the relationship between chromosomal architecture and plasmids across 52,393 complete prokaryotic genomes. We demonstrate that chromosomal gene content robustly predicts multiple dimensions of plasmid biology, including carriage status, quantitative load, mobility capabilities and functional cargo composition. Remarkably, predictive signals can be compressed to as few as 15 genes—or simply the global composition of seven major enzyme classes. Together, these findings support a conceptual framework in which microbial chromosomes encode an intrinsic constraint on plasmids, establishing chromosomal architecture as a determinant of evolutionary potential rather than merely the genetic background upon which HGT acts.

## Results

### Model development and evaluation

To systematically investigate the predictability of plasmid carriage, we assembled a dataset of 52,393 complete prokaryotic genomes from RefSeq^25^ (release of August 1, 2025), spanning diverse bacterial and archaeal lineages (Fig. 1b). All genomes were uniformly annotated with the NCBI Prokaryotic Genome Annotation Pipeline (PGAP)^26^. Plasmid carriage varied substantially across genomes (Fig. S1a): the average plasmid count per genome was 1.22, with 45.8% of genomes harboring at least one plasmid (Fig. S1b). Plasmid prevalence differed markedly among taxa and even among strains of the same species. In *Escherichia coli*, for instance, plasmid numbers ranged from 0 to 18 per genome, with 15.32% of genomes lacking plasmids entirely. This variation provides a foundation for linking host chromosomal features to plasmid carriage.

From all the chromosomes, 72,451 non-redundant genes are identified (Fig. S1c), each with a unique product descriptor (e.g., ‘amino acid permease’ or ‘16S ribosomal RNA’). To identify chromosomal features predictive of plasmid carriage, we first evaluated each gene’s association with plasmid presence using chi-square tests with Benjamini–Hochberg FDR correction and log₂ fold change filtering. After removing phylogenetically confounded signals (genes restricted to a single genus or present in <1% of genomes), we retained a core set of 2,824 chromosomal genes (Fig. 1c, see Methods for more details). For each genome, we calculated the abundance of these genes on the chromosome and used them as predictive features (Fig. 1d).

We first asked whether this feature set could predict plasmid presence—a binary classification problem. A random forest (RF) classifier achieved a mean AUROC (hereafter referred to as AUC) of 0.919 ± 0.002, an accuracy of 0.843 ± 0.003, and an AUPRC of 0.907 ± 0.003 across five independent train–test splits (Fig. S2), suggesting that plasmid presence is highly predictable from host chromosomal gene content. Notably, most false negatives carried only minimal plasmid content, indicating that errors arise mainly from genomes with marginal plasmid signal (Fig. S3). Benchmarking against alternative algorithms, RF consistently outperformed *k*-NN (*k*=1,2,…,10) (Fig. S4), Extra Trees, Logistic Regression, Naive Bayes, and Histogram-based Gradient Boosted Trees (Fig. 1e, see Methods for more details).

Beyond binary prediction, we extended the framework to quantitative aspects of plasmid load. A RF regressor trained on the same feature set predicted plasmid number per genome with reliable accuracy (R² = 0.584±0.012) (Fig. 1f). RF-based prediction of total plasmid size—a continuous measure summing all plasmids in a genome—achieved even higher performance (R² = 0.773±0.018), outperforming alternative regressors (Fig. 1g and h).

Because plasmid carriage varies across taxonomic lineages, we next asked whether predictive performance merely reflected phylogenetic structure. Stratifying the test genomes by phylum revealed consistently high AUCs for plasmid carriage classification within each major prokaryotic group (Fig. 1i, S5). Performance of plasmid size regression varied more across phyla, largely due to uneven genome numbers, but remained reliable in most phyla. To further minimize phylogenetic effects, we evaluated the test performance within the two well-represented genera, *Escherichia* and *Salmonella*. Despite the substantially reduced genomic variation within each genus, both plasmid carriage and size remained highly predictable (Fig. S6), indicating that the predictive signal persists even among closely related genomes. These results suggest that the model captures predictive features of plasmid load beyond coarse taxonomic identity.

Chromosome size is another known correlate of plasmid carriage^27,28^. To determine whether model performance was driven primarily by chromosomal size, we partitioned test genomes into bins by chromosome length, minimizing size effects within each bin. Prediction accuracy remained high across most bins (Fig. S7a-c). As a complementary control, we converted gene counts to chromosome-size-normalized density (copies per kb) and trained a RF regressor to predict total plasmid size directly from this density-based feature set, which still achieved R² = 0.745 ± 0.018, confirming that predictive signal persists even after explicitly removing the chromosome-size confound (Fig. S7d). Together, these results suggest that the predictive power is not solely attributable to phylogenetic relatedness or chromosome size, but reflects intrinsic features of chromosomal genetic architecture.

We next asked whether chromosomal content could predict plasmid mobility—a key driver of HGT dynamics. Integrating mobility annotations for all plasmids (genomes harboring multiple plasmids were assigned the highest mobility class among the plasmids)^29^, we trained a three-class RF classifier (non-mobilizable, mobilizable, conjugative) using the one-vs-rest strategy (Fig. 1j). The model achieved a mean macro-averaged AUC of 0.907 and a mean macro-averaged AUPRC of 0.831 across independent replicates, with per-class AUC values exceeding 0.87 for all categories (Fig. 1k and Fig. S8). Together, these results demonstrate that the chromosomal gene content encodes a predictive signature for plasmid mobility potential.

### Robustness and generalizability of the predictive models

To assess the robustness of the predictions against data noise, we subjected the RF models to a series of controlled challenges. First, simulating chromosome incompleteness typical of draft genomes or metagenomic assemblies, we randomly removed fractions of gene records from each chromosome in the test set. For plasmid presence classification, the AUC declined from 0.919 (no removal) to 0.897 (10% removal) (Fig. S9a). The size regressor was more sensitive, but retained an R² of 0.577 at 10% removal (Fig. S9b). These results suggest that binary classification withstands substantial gene loss, while quantitative size prediction requires more complete genomic context.

Second, the boundary between plasmids and chromosomes can be ambiguous—small secondary chromosomes may be misannotated as plasmids, or megaplasmids as chromosomes, leading to labeling errors of plasmid carriage status^30^. To evaluate robustness against such errors, we randomly flipped plasmid presence labels in the training set while keeping the test set unchanged. The model maintained an AUC > 0.86 even with 20% label flipping (Fig. S9c), demonstrating substantial tolerance to mislabeled instances.

Third, we evaluated resilience to inconsistencies in gene product nomenclature. While RefSeq genomes were annotated under a standardized pipeline that maximized naming consistency, rare cases exist where the same gene is recorded under distinct names across taxa, causing it to be treated as multiple distinct features. To simulate such redundancy, we randomly merged 1%–95% of feature columns. The AUC remained stable even when 80% of features were collapsed into a single dimension (Fig. S9d); the size regression model showed comparable robustness (Fig. S9e). We further repeated the prediction pipeline using another strategy to determine gene identity, by mapping gene product names to UniProt Pfam domain identifiers^31^. This reduced the feature space to 1,710 dimensions without compromising performance (Fig. S9f). The size regression model showed the same pattern, achieving an R² of 0.773 under the Pfam-derived features (Fig. S9g). Together, these results demonstrate that the model learns from biologically meaningful genomic architecture rather than naming idiosyncrasies.

To test whether the models generalize to novel lineages rather than memorizing species-specific signatures, we implemented two complementary strategies. First (species de-replication), we collapsed the dataset to one representative genome (randomly selected) per species (n = 7,991) and performed a random split. The model achieved a test AUC of 0.796 (Fig. S10a), confirming substantial predictive power even after removing within-species redundancy. The size regression model yielded an R² of 0.553 (Fig. S10b), reflecting the reduced sample size and the greater challenge of cross-species quantitative prediction. Second (species holdout), we reserved 20% of species for testing and excluded their genomes from training, evaluating the model exclusively on unseen species. The model achieved a test AUC of 0.815 and an accuracy of 0.754 (Fig. S10c). Under this scheme, the size regression model achieved an R² of 0.638 (Fig. S10d). Together, these results demonstrate that the predictive signal transcends phylogenetic boundaries and reflects a generalizable biological relationship.

Finally, we assessed temporal generalizability using an RefSeq dataset released after model development, comprising 10,784 bacterial genomes (5,396 plasmid-carrying genomes and 15,278 plasmids). Without further feature selection or hyperparameter adjustment, the models retained high predictive performance, achieving an R² of 0.799 for plasmid-size prediction and an AUC of 0.889 for plasmid-carriage classification (Fig. S11). These results demonstrate that plasmid compatibility is not a transient property inferred from historical genome collections, but a stable genomic feature that persists across newly documented bacterial diversity.

### Derivation of minimal predictive signatures and functional determinants

To determine how predictive information is distributed across chromosomal features, we first ranked all 2,824 genes according to their mean decrease in Gini impurity (MDI) importance, averaged across the five random forest refits^32^. Predictive contributions were highly concentrated: the top 10 genes accounted for 10.7% of the total importance, and half of the cumulative importance was captured by only 263 genes (9.3% of all features), indicating that the predictive signal for plasmid carriage is concentrated within a relatively small fraction of the feature set.

We next asked how far this feature space could be compressed without substantially compromising predictive performance. Incremental feature selection revealed that classification performance rapidly plateaued with only 15 genes (Fig. 2a), representing a >99% dimensionality reduction relative to the full model. Despite its simplicity, this minimal signature generalized across multiple dimensions of plasmid biology. Models trained exclusively on these 15 genes predicted total plasmid size with an R² of 0.669 (Fig. 2b) and genome-level plasmid mobility with a macro-averaged AUC of 0.893 (Fig. 2c), approaching the performance of the complete feature set. Thus, diverse plasmid-associated traits can be inferred from a remarkably compact chromosomal signature.

**Figure 2.**
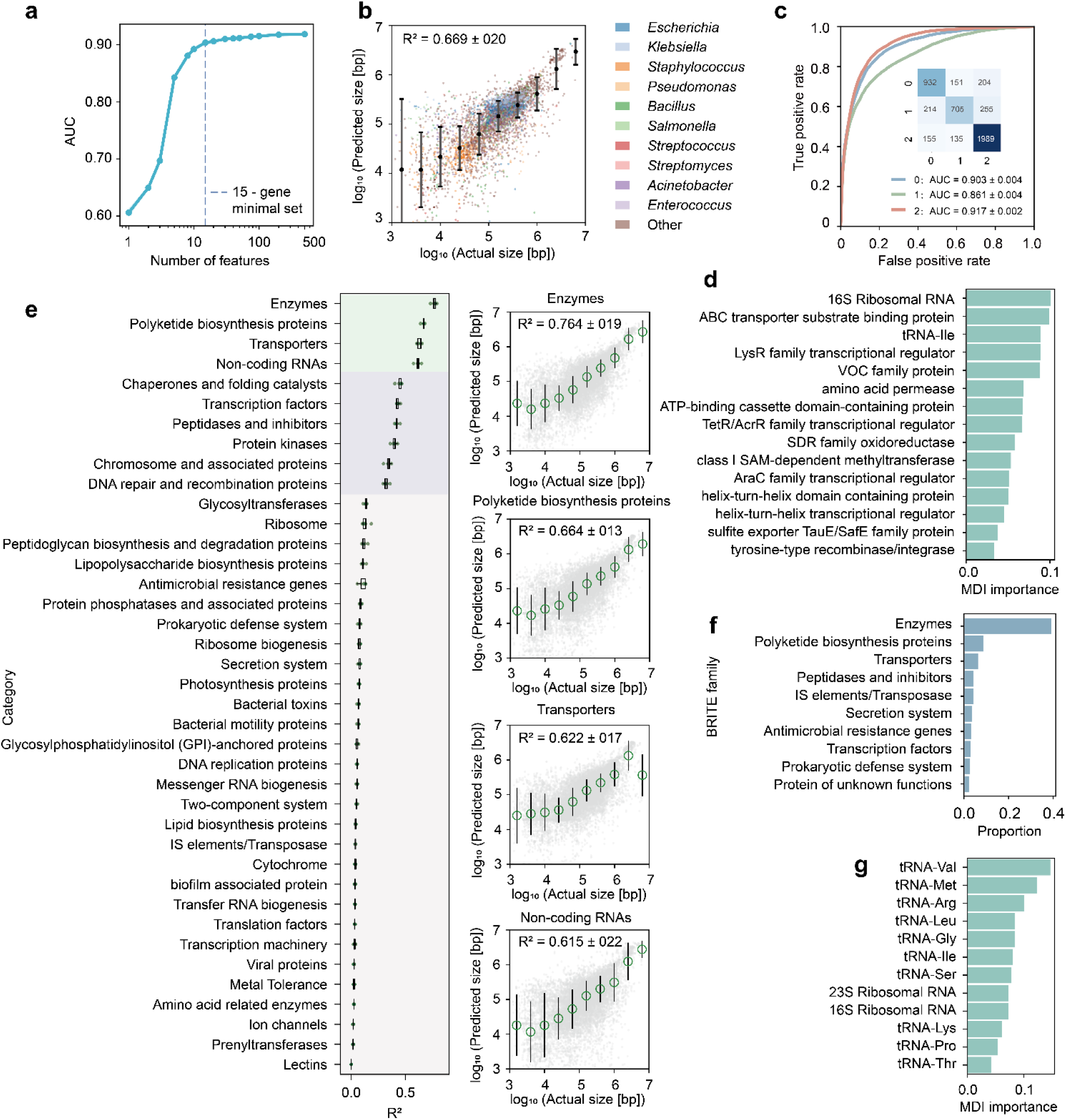
Compact predictive signatures and functional determinants of plasmid biology. (a) Incremental feature selection (IFS) curve for the plasmid carriage classifier. Features were ranked by mean Gini importance. The solid line and shaded band represent the mean AUC ± s.d. on the test sets of five replicates. The dashed vertical line marks the minimal predictor set (k = 15), defined as the smallest feature subset achieving ≥ 98% of the maximum observed AUC. (b) Prediction of total plasmid size using the 15-gene minimal feature set. Points represent individual genomes from a test set and are colored by the ten most abundant genera. Both axes are displayed on a log₁₀ scale (bp). Filled circles indicate mean predicted values within bins of observed plasmid size, and error bars denote ± s.d. of predicted values within each bin. The reported R² value corresponds to the mean ± s.d. across five independent train–test splits. (c) Prediction of genome-level plasmid mobility using the 15-gene product set. Mobility classes are defined as non-mobilizable (0), mobilizable (1), and conjugative (2). ROC curves show the mean one-versus-rest performance across five independent train–test splits, with per-class AUC values reported as mean ± s.d across the five independent train–test splits. (d) Mean decrease in impurity (MDI) scores of the 15-gene product set for plasmid size prediction. Bars represent mean MDI scores derived from five independent Random Forest regressors and normalized to sum to 1. Features are ordered by decreasing importance. (e) Predictive power of separate KEGG BRITE functional categories for plasmid-size prediction. Left, distribution of test-set R² values for 39 functional categories evaluated using category-specific Random Forest regressors. Boxplots show the median (center line), interquartile range (box limits), and minimum–maximum values (whiskers), with 5 individual replicate scores overlaid as dots. Right, observed versus predicted plasmid size for the four highest-performing categories. Points represent pooled predictions from all five train–test splits. Both axes are shown on a log₁₀ scale (bp). Filled circles indicate mean predicted values within bins of observed plasmid size, and error bars denote ± s.d. The reported R² values represent mean ± s.d. across 5 replicates. (f) Proportion of ten most abundant functional categories within the feature gene set. (g) MDI scores of the 12 non-coding RNA features for plasmid size prediction. Bars represent mean MDI scores across five independent Random Forest regressors, normalized to sum to 1 and ranked by decreasing importance.

The minimal predictor set was dominated by regulators and genome-plasticity functions, including helix–turn–helix transcription factors, LysR-and TetR/AcrR-family regulators, tyrosine recombinases/integrases, class I SAM-dependent methyltransferases, and ATP-binding cassette proteins (Fig. 2d). These genes showed pronounced phylogenetic variation^33^, with the highest abundances in *Pseudomonadota* and *Actinomycetota* (Fig. S12).

We next sought to determine which biological functions contribute most to the predictive signal. Chromosomal features were grouped into 39 KEGG BRITE functional categories^34^, and models were retrained independently using each category (see Methods for more details). Predictive performance varied substantially among functional groups, with enzyme-encoding genes emerging as the strongest predictive features (AUC = 0.911), followed by polyketide biosynthesis proteins and transporters (Fig. 2e and f). Unexpectedly, non-coding RNAs ranked among the most informative categories despite comprising only 12 features. This compact set, consisting of two ribosomal RNAs and ten transfer RNAs (Fig. 2g and Fig. S13), achieved an AUC of 0.859 for plasmid carriage prediction and an R² of 0.615 for plasmid size regression, demonstrating that highly compressed translational signatures retain substantial predictive power.

Collectively, these analyses reveal two complementary principles underlying chromosomal prediction of plasmid biology. First, predictive information is highly compressible, with a small set of genes sufficient to recover most of the model performance. Second, this information is functionally structured rather than uniformly distributed, with enzyme-associated genes representing the dominant source of predictive power. This observation motivated our subsequent analysis of host metabolism as a systems-level determinant of plasmid compatibility.

### Enzymatic organization encodes plasmid compatibility

Enzyme-encoding genes were the strongest functional predictive features for plasmid prediction, raising a fundamental question: does this predictive signal originate from a small number of individual enzymes, or from the higher-level metabolic organization? To distinguish between these possibilities, we projected enzyme-associated genes onto the hierarchical Enzyme Commission (EC) framework^35^, progressively compressing the feature space from 2,622 specific enzymatic reactions (EC4) to only seven major classes (EC1) (Fig. 3a).

**Figure 3.**
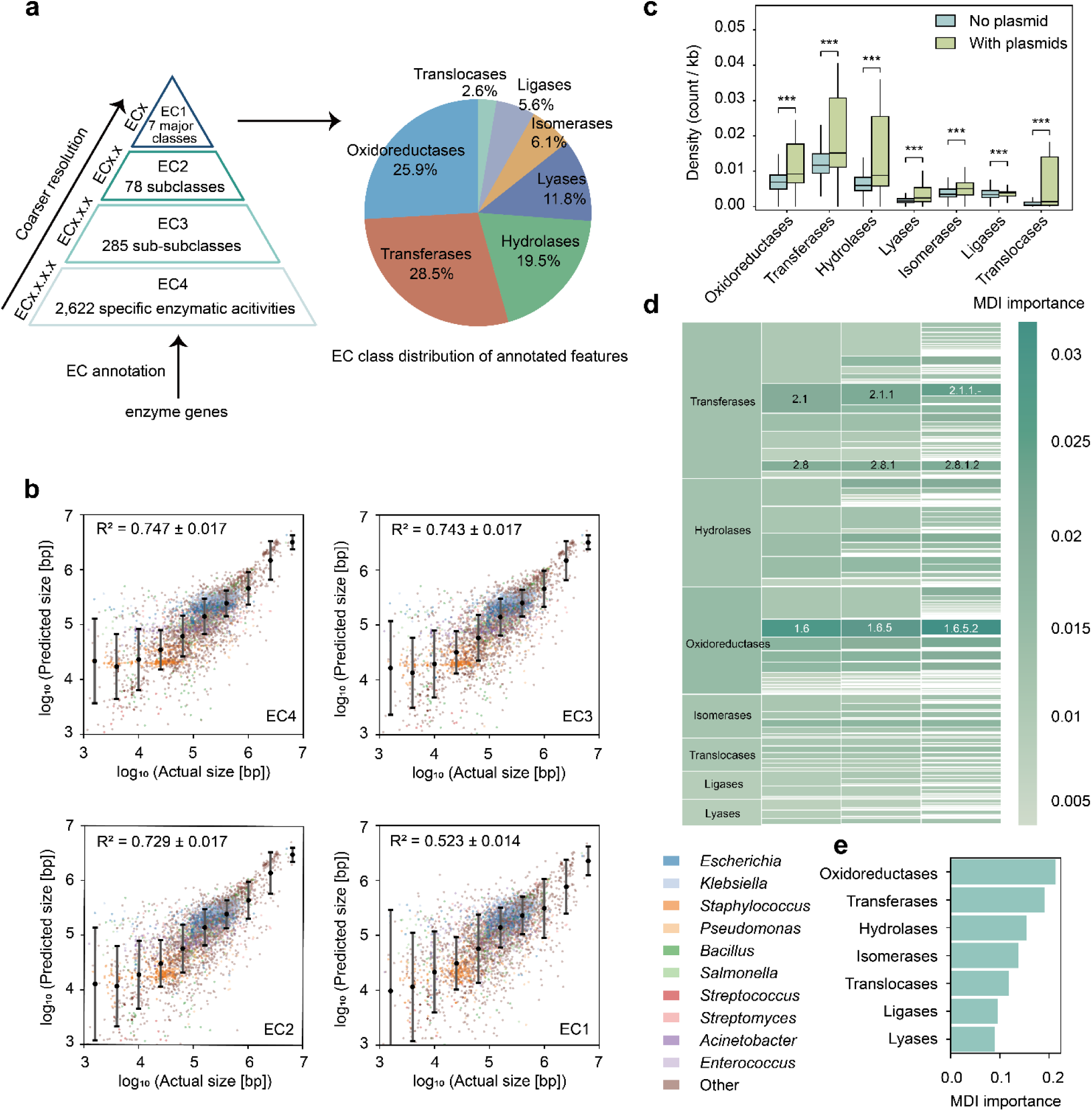
Enzymatic architecture predicts plasmid loads and reveals metabolic restructuring associated with plasmid carriage. (a) Construction of hierarchical Enzyme Commission (EC) feature representations. Chromosomal enzyme-associated genes were identified from RefSeq product annotations and mapped to EC identifiers. Genome-level EC abundance profiles were generated by summing the chromosomal copy numbers of genes assigned to the same EC entry. Hierarchical EC feature representations were then constructed by progressively aggregating EC annotations from EC4 (2,622 specific enzymatic reactions) to EC3 (285 sub-subclasses), EC2 (78 subclasses), and EC1 (7 major enzyme classes), providing functional representations at multiple levels of enzymatic resolution. The pie chart summarizes the distribution of annotated features across the seven EC1 enzyme classes, including transferases, oxidoreductases, hydrolases, lyases, isomerases, ligases, and translocases. (b) Performance of plasmid size regression across four EC levels. Scatter plots show observed versus predicted plasmid sizes (log₁₀ bp) for a representative test partition. Points represent individual genomes and are colored by the ten most abundant genera. Filled circles indicate mean predicted values within bins of observed plasmid size; error bars denote ± s.d. The reported R² values represent mean ± s.d. across five independent train–test splits. (c) Global differences in chromosomal enzyme composition associated with plasmid carriage. Enzyme density was calculated as the number of chromosome-encoded enzymes assigned to each of the seven EC1 classes per kilobase of chromosomal sequence and compared between plasmid-free and plasmid-bearing genomes. Boxes indicate the median and interquartile range, and whiskers extend to 1.5 times the interquartile range; outliers are not shown. Statistical significance was assessed using two-sided Mann–Whitney U tests, with Benjamini–Hochberg false-discovery-rate correction across the seven enzyme classes. Three asterisks represent *q* value below 0.001. (d) Hierarchical distribution of mean decrease in impurity (MDI) scores across EC levels for plasmid-size prediction. At the EC4 level, rectangle area and color intensity represent individual MDI scores. At higher EC levels, rectangle area represents cumulative MDI scores aggregated from descendant EC4 features, whereas color intensity represents the MDI-weighted mean of the corresponding descendant features. (e) Relative importance of the seven top-level EC classes for plasmid size prediction. Horizontal bars show mean MDI scores ± s.d. across five independent Random Forest regressors, normalized to sum to 1.

Despite the drastic reduction in feature dimensionality, predictive performance remained remarkably high. Models trained using only the seven EC1 classes achieved an AUC of 0.874 for plasmid carriage and an R² of 0.523 for total plasmid size, while progressively finer EC resolutions gradually restored predictive performance, with EC4 approaching that of the original gene-level model (Fig. 3b, Fig. S14). Thus, the predictive signal is not concentrated in a limited set of enzymes, but is broadly distributed across the catalytic architecture of the chromosome.

Having established that predictive information is distributed across metabolism, we next asked whether it depends on absolute enzyme abundance or on the relative organization of metabolic functions. Plasmid-carrying genomes displayed coordinated shifts across all seven EC1 classes, irrespective of whether enzyme repertoires were quantified by absolute abundance or genomic density (Fig. 3c). These shifts were also evident within *Escherichia* and *Salmonella* (Fig. S15), indicating that the association was retained within individual genera. A model trained solely on the relative proportions of the seven EC1 classes achieved an AUC of 0.868 for plasmid carriage and an R² of 0.447 for plasmid size. Predictive capacity was albeit reduced within *Escherichia* and *Salmonella* (Fig. S16). Principal component analysis (PCA) of the seven EC1 features further characterized the major axes of variation in chromosomal enzymatic architecture. Mapping log₁₀-transformed total plasmid size onto this PCA space revealed its distribution across the principal components (Fig. S17a). These findings indicate that plasmid compatibility is encoded largely in the relative allocation of catalytic functions, independent of genome size or absolute enzyme abundance.

We next asked which metabolic functions contribute most strongly to this distributed signal. At the finest resolution, transferases were the dominant predictive features of both plasmid carriage and plasmid size (Fig. 3d, Fig. S17b). These importance patterns were highly stable across hierarchical resolutions: aggregating EC4 feature importance to EC3 closely reproduced those learned directly from EC3 models (Pearson’s *r* = 0.902, *p*-value < 0.001; Fig. S17c). At the coarsest level, however, the dominant contributors diverged, with oxidoreductases contributing most strongly to plasmid size, whereas hydrolases contributed most to plasmid carriage (Fig. 3e, Fig. S17d). To assess whether model-derived importance reflected marginal associations, we compared EC1 MDI scores with Pearson correlations to untransformed total plasmid size. MDI importance showed strong concordance with feature–target correlation across the seven EC1 classes (Pearson’s *r* = 0.835, *p*-value = 0.019; Spearman’s *ρ* = 0.857, *p*-value = 0.014; Fig. S17e), indicating that EC1 classes more positively associated with plasmid size generally received greater importance in the RF model. Together, these results reveal that plasmid compatibility is encoded hierarchically within host metabolism, with distinct metabolic layers constraining different dimensions of plasmid biology.

### Chromosomal architecture predicts plasmid functional payloads

Having shown that chromosomal architecture predicts plasmid carriage, abundance and mobility, we next asked whether it also predicts the functional cargo encoded by plasmids. Because antibiotic resistance represents one of the most consequential plasmid-associated traits, we first quantified plasmid-borne antibiotic resistance genes (ARGs) across 33 resistance classes and evaluated their predictability from host chromosomal features^36^.

Multiple clinically important resistance classes were predicted with substantial accuracy, including glycopeptide (R² = 0.624), cephalosporin (R² = 0.605), penicillin β-lactam (R² = 0.573) and monobactam resistance (R² = 0.537) (Fig. 4a, b). Across genomes with the highest plasmid ARG burdens, predicted resistance profiles closely recapitulated the observed multidrug resistance landscapes (Fig. 4c), indicating that host chromosomal composition contains substantial information about the functional repertoire of resident plasmids.

**Figure 4.**
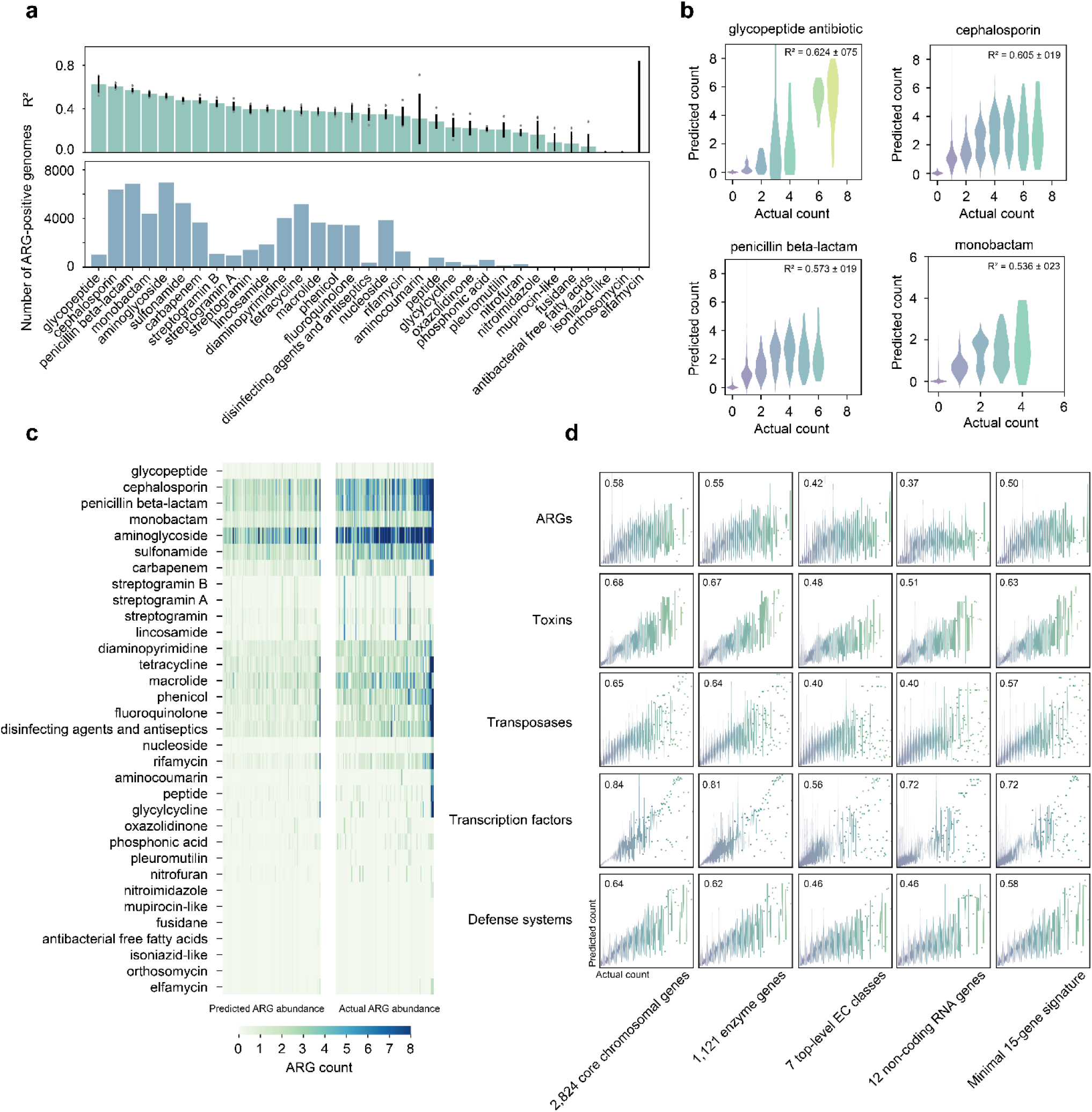
Host chromosomal architecture predicts plasmid-encoded functional repertoires. (a) Predictive performance across 33 plasmid-encoded antibiotic resistance gene (ARG) classes. Random Forest regressors were trained using chromosomal features from plasmid-carrying genomes and evaluated across five independent train-test splits. ARG classes are ranked by mean test-set R². Bars and error bars indicate mean ± s.d., with individual replicate values overlaid as dots. Bottom, number of genomes carrying at least one ARG on plasmids for each resistance class. (b) Prediction of the four best-predicted ARG classes. Bean plots show the distribution of predicted counts for each observed count category in a representative test partition. Bean width represents the local kernel density estimate. R² value represents mean ± s.d. across five independent train–test splits. (c) Prediction of plasmid-borne ARG abundance from chromosomal gene content. Heatmaps compare Random Forest–predicted (left) and observed (right) abundances of 33 plasmid-associated ARG classes in plasmid-carrying genomes with the highest total ARG burden. Columns represent individual test genomes (top 2% by total plasmid-borne ARG abundance; n = 96), ordered from left to right by increasing total ARG burden. Rows denote ARG classes, and color intensity indicates class-specific copy number per genome. (d) Prediction of major plasmid functional cargo across five chromosomal feature representations. Rows correspond to five plasmid functional categories: ARGs, toxins, transposases, transcription factors, and defense systems. Columns represent five chromosomal feature sets: 2,824 feature genes; 1,121 enzyme genes; seven EC1 enzyme classes; 12 non-coding RNAs; and the 15-gene minimal predictor set. Each panel shows observed versus predicted values, with the corresponding test-set R² indicated in the upper-left corner.

We next asked whether this predictive relationship extends beyond antibiotic resistance. Using the same chromosomal framework, we accurately predicted the plasmid-encoded abundance of toxins, transposases, transcription factors and prokaryotic defense systems (Fig. 4d; see Methods for details). This predictive capacity was consistently retained across multiple chromosomal feature representations, including enzyme-associated genes, EC enzyme classes, non-coding RNAs and the parsimonious 15-gene signature, demonstrating that functional payload prediction is not dependent on a particular feature space. Predictive performance, albeit reduced, was also observed within individual genera, including *Escherichia* and *Salmonella* (Fig. S18).

Together, these results extend chromosomal prediction from plasmid burden to plasmid function, demonstrating that host genomes encode a latent blueprint not only of whether plasmids can be maintained, but also of the mobile genetic functions they can accommodate.

## Discussion

Horizontal gene transfer is commonly viewed as a process governed by plasmid mobility and environmental selection^1,37,38^. In this framework, the chromosome is largely regarded as the genomic context in which mobile elements operate. Our results suggest a different perspective. Across more than 52,000 genomes, diverse properties of plasmids—including their presence, abundance, mobility and functional repertoire—were predictable from the host chromosomes. This implies that bacterial chromosomes are not passive substrates for horizontal gene transfer, but contain genomic signatures associated with the capacity of hosts to accommodate different plasmid types^39^. Rather than representing a fixed property imposed by individual genes, plasmid compatibility appears as an emergent property of host genomic organization, defining an evolutionary landscape within which mobile elements circulate.

An important consideration is that plasmid–chromosome relationships are inherently bidirectional: plasmids can reshape chromosomes through gene acquisition, sequence exchange and long-term coevolution^40,41^, and chromosomal features associated with plasmid carriage may therefore represent both determinants and historical signatures of this interaction. Our results do not distinguish these evolutionary directions; instead, they reveal that the genomic state of the host contains sufficient information to predict its present-day plasmid compatibility landscape. This predictive relationship is consistent with a model in which chromosome and plasmid evolution are coupled^23,42^, with host genomic architecture both reflecting past mobile-element interactions and constraining future trajectories of horizontal gene transfer.

Our findings provide an opportunity to reconcile two prevailing views of plasmid ecology. Environmental selection undoubtedly determines which plasmids are advantageous under a given condition^43^, yet selection operates within the range of host–plasmid combinations that are physiologically and evolutionarily feasible^27,44^. Chromosomal architecture therefore defines the feasible space upon which environmental selection acts^45,46^. Rather than representing competing explanations, host genomic architecture and environmental selection act at different hierarchical levels: the former establishes the boundaries of compatibility, whereas the latter determines which compatible plasmids are ultimately favored.

A striking observation is that predictive information persists despite successive compression of the feature space—from thousands of genes to seven enzyme classes, and even to a small set of translational RNAs. This hierarchical robustness argues against models in which plasmid compatibility is determined by a limited number of molecular interactions. Instead, compatibility emerges as a systems-level property of the cell, arising from the integrated organization of metabolism and resource allocation rather than individual genes. The host genome therefore appears to encode a distributed representation of its capacity to support mobile genetic elements.

The ability to predict plasmid functional cargo from chromosomal architecture suggests that horizontal gene transfer is not an unconstrained process of gene movement^17,47–49^. Instead, different mobile functions occupy different regions of the host compatibility landscape. Antibiotic resistance genes^36^ and defense systems^50–55^ therefore disseminate through bacterial populations under distinct host-dependent constraints, providing a mechanistic explanation for why some resistance determinants repeatedly emerge within particular bacterial lineages whereas others do not^38^.

Our framework also provides implications for predictive microbial genomics. In microbial ecology, compatibility landscapes may improve predictions of plasmid dissemination across natural communities and enable digital reconstruction of horizontal gene transfer potential from genomic surveys alone. In antimicrobial resistance research, host chromosomal determinants associated with plasmid persistence may represent targets for destabilizing resistance plasmids independently of conventional antibiotics^56–58^. In synthetic biology, the compact chromosomal signatures identified here may provide a rational basis for selecting or engineering microbial chassis with enhanced plasmid stability and predictable maintenance. Because the framework relies exclusively on chromosomal information, it is readily applicable to newly sequenced genomes without prior knowledge of their plasmid content.

Several limitations should be acknowledged. Our analyses were restricted to complete RefSeq genomes and therefore incompletely represent the diversity of uncultivated microorganisms recovered from metagenomic assemblies^59^. Furthermore, the relationships identified here are predictive rather than causal. Determining how metabolic organization, translational capacity and other host properties mechanistically influence plasmid compatibility will require targeted experimental perturbation and long-term evolution experiments. Extending this framework to other classes of mobile genetic elements—including bacteriophages, integrative conjugative elements and genomic islands—will further test whether compatibility landscapes represent a general organizing principle of microbial genome evolution^60,61^.

Ultimately, our study indicates that chromosomes do not merely provide the genetic background in which plasmids evolve—they define the evolutionary landscape on which plasmid evolution occurs.

By encoding the intrinsic capacity of a host to accommodate mobile DNA, genome architecture places predictable constraints on horizontal gene transfer, plasmid persistence and the dissemination of adaptive traits. Viewing chromosomes as determinants of evolutionary potential rather than passive genetic substrates provides a conceptual framework for predicting, and ultimately engineering, the flow of genetic information across microbial communities.

## Methods

### Data curation

All prokaryotic genomes were retrieved from the NCBI RefSeq database (release of August 1, 2025). To ensure consistent genome quality and annotation standards, only assemblies classified as complete genome, annotated by RefSeq, and not designated as atypical were included. Genome records were downloaded programmatically using the NCBI Datasets command-line tool (v2) as GenBank flat files^62^, which contain both nucleotide sequences and gene annotations generated by the NCBI Prokaryotic Genome Annotation Pipeline (PGAP). Assemblies that could not be successfully retrieved were excluded from further analysis. The final dataset comprised 52,393 complete bacterial and archaeal genomes.

GenBank flat files were parsed using custom Python scripts to extract plasmid and chromosomal features. Replicons annotated as plasmids were identified from RefSeq records and used to calculate plasmid number and total plasmid size for each genome. For assemblies containing multiple chromosomes, all chromosomal replicons were merged and treated as a single chromosomal gene set. Gene identities were defined by their PGAP product descriptors. Entries annotated as “hypothetical protein” were excluded, yielding 72,451 non-redundant gene descriptors across the dataset. For each genome, the copy number of every gene descriptor was quantified across the aggregated chromosomal replicons, generating a genome-by-gene abundance matrix that was used for all downstream analyses and predictive modelling.

### Feature extraction

For each chromosomal gene, we quantified its association with plasmid carriage across all genomes. Let *x_i_* denote the copy number of a given gene in the chromosomes of the *i*-th genome, and *y_i_* a binary variable indicating plasmid carriage status, with *y_i_* = 1 if the genome carries at least one plasmid and *y_i_* = 0 otherwise. For statistical testing, gene abundance was binarized to gene presence or absence. Associations between gene occurrence and plasmid carriage were evaluated using Pearson’s chi-square test, and resulting *P* values were adjusted using the Benjamini-Hochberg false discovery rate (FDR) procedure^63^.

To quantify the magnitude and direction of the association, we calculated the log_2_ fold change (log_2_*FC*) in plasmid-carriage prevalence between genomes containing and lacking the target gene on chromosomes:

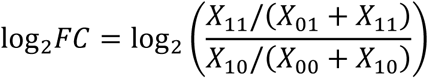

where *X*_11_ and *X*_01_ represent the number of plasmid-carrying and plasmid-free genomes that encode the target gene on chromosomes, respectively, and *X*_10_ and *X*_00_ denote the corresponding counts for genomes lacking the gene on chromosomes.

Genes with Benjamini-Hochberg FDR-adjusted *p*-value < 0.05 and | log2*FC* | > 1.0 were retained, yielding 12,262 candidate features. To ensure broad taxonomic relevance and generalizability, genes restricted to a single genus were removed, reducing the feature set to 7,625. A second prevalence filter excluded genes present in fewer than 1% of genomes (< 524 genomes), resulting in a final set of 2,824 chromosomal features. For each genome, the chromosomal copy number of these genes were compiled to a feature vector [*N*_1_, *N*_2_,…, *N_m_*], where *N_j_* represents the chromosomal abundance of the *j*-th feature. These vectors served as inputs for downstream predictive modeling.

### Model development

To predict plasmid carriage and plasmid loads from chromosomal gene content, we constructed supervised machine-learning models using the abundance profiles of the 2,824 selected chromosomal genes as input features. Genomes were randomly partitioned into training (80%) and test (20%) sets, and model performance was evaluated using five independent random splits. For plasmid carriage, we implemented a binary classification framework, whereas total plasmid size and plasmid number were treated as regression tasks.

For total plasmid size regression, we compared multiple machine learning algorithms including Random Forest (RF)^64^, Extra Trees^65^, Ridge Regression (coupled with standard scaling)^66^, Bayesian Ridge^67^, and Histogram-based Gradient Boosting (Hist GBDT)^68^. For plasmid carriage classification, we compared RF, Extra Trees, Logistic Regression (with standard scaling)^69^, Naïve Bayes^70^, and Hist GBDT. In addition, *k*-nearest-neighbor (*k*-NN) models (*k*=1, 3, 5, 7, 10) were evaluated as similarity-based baselines to determine whether plasmid-associated signals could be explained primarily by overall genomic resemblance^71^. Across all prediction tasks, RF consistently achieved the highest performance and was therefore selected for subsequent analyses.

To maximize predictive capacity, hyperparameters for the RF-based size regressor and presence classifier were independently optimized via sequential random and grid search. The final size regressor was configured with 200 decision trees, maximum features set to the square root, minimum samples per split = 5, minimum samples per leaf = 1, maximum tree depth constrained to 30 layers (to prevent overfitting on the wide-range continuous target), and bootstrap sampling disabled. The classifier was configured with 110 unconstrained trees, log₂ maximum features, minimum samples per split = 6, minimum samples per leaf = 1, and bootstrap disabled. We extended this RF-based framework to the prediction of plasmid number using a standard RF regressor with the same hyperparameter configuration as the size model.

Classification performance was evaluated using the AUC, AUPRC, accuracy, precision, recall and F1 score. Regression performance was assessed using the coefficient of determination (R²). To assess the robustness of the predictive framework across hosts with different chromosome sizes, test genomes were stratified by chromosome size, and predictive performance was quantified separately within each size bin.

To evaluate whether predictive performance was driven by lineage-specific signals, we assessed model performance across taxonomic boundaries using a post-hoc stratification strategy. Models were trained on randomly partitioned dataset (80% training, 20% test) without imposing any taxonomic constraints during training. After prediction, test genomes were assigned taxonomic annotations using TaxonKit based on the NCBI Taxonomy database^72,73^, enabling performance to be quantified within taxonomic groups.

To ensure robustness, analyses were restricted to phyla represented by at least 100 genomes in the test set. Standard classification metrics, including true positives (TP), false positives (FP), and false negatives (FN) were determined within each phylum. For regression tasks, performance was quantified as the coefficient of determination (R²) between predicted and observed plasmid sizes within each phylum. To further assess genus-level robustness, we separately evaluated predictions for *Escherichia* (n = 4,203; plasmid prevalence 84.7%) and *Salmonella* (n = 1,853; 70.1%) from the test set.

### Prediction of plasmid mobility

Plasmid mobility was annotated using MOB-suite^29^, which categorizes plasmids as conjugative, mobilizable, or non-mobilizable based on the presence of relaxase and type IV secretion system (T4SS) components. Because individual genomes frequently harbor multiple plasmids with distinct mobility types, we derived a single genome-level mobility label by assigning each genome the highest mobility class observed among its constituent plasmids. Genomes lacking mobility annotations were excluded, yielding a dataset of 23,717 genomes comprising conjugative (48.1%), mobilizable (24.8%) and non-mobilizable categories.

To predict plasmid mobility from host genomic architecture, we trained a multiclass Random Forest classifier using the abundance profiles of the 2,824 chromosomal signature genes as input features. Model performance was evaluated across five independent stratified 80:20 train–test splits. Classification performance was evaluated using the AUC under a one-versus-rest framework, with performance assessed separately for each mobility class and subsequently averaged across classes.

## Robustness validation

To determine whether predictive performance reflects robust biological signal rather than dataset-specific artifacts, we systematically evaluated model stability under multiple perturbation regimes.

(1) Genomic completeness sensitivity. To simulate incomplete genome assemblies, a gene dropout procedure was applied in which 80%–100% of chromosomal gene entries were randomly subsampled without replacement at 5% intervals. Models were trained on complete genomes and evaluated on corresponding degraded test inputs, thereby isolating the effect of input incompleteness on predictive performance.
(2) Label noise robustness. To assess resilience against label annotation errors—such as ambiguous boundary calls between megaplasmids and secondary chromosomes, stochastic label noise was introduced exclusively into the training set by randomly flipping 1%–20% of class labels (plasmid presence↔absence). Evaluation was performed on an unperturbed test set to quantify degradation under controlled label corruption.
(3) Feature space perturbation and redundancy collapse. To test sensitivity to gene-level naming redundancy and nomenclature inconsistency, subsets of features (1%–95%) were randomly selected and collapsed into composite features via summation, applied consistently to both training and test sets.
(4) Functional re-annotation using protein families. To further evaluate whether predictive power depends on gene nomenclature, the feature space was re-encoded at the level of Pfam domains. The 2,824 gene products were mapped to Pfam identifiers via UniProt REST API^31,74^, and gene counts sharing identical domains were aggregated within each genome. After exclusion of features lacking Pfam assignments, a 1,710-dimensional domain-level representation was constructed.
(5) Phylogenetic redundancy control. To quantify the contribution of taxonomic structure, two strategies were implemented. First, a species de-replication approach retained only one genome per species prior to partitioning. Second, a stringent species holdout strategy excluded all genomes from 20% of species during training. The model was then evaluated on these unseen lineages.
(6) To evaluate generalizability beyond the 52,393 genomes used for training, we constructed an independent test set of 10,784 complete bacterial and archaeal genomes from a later RefSeq release (post-August 1, 2025). Chromosomal gene-product abundance was quantified using the same annotation and feature-extraction pipeline, yielding the same set of 2,824 chromosomal gene-product features. The final models, trained on the complete training dataset, were directly applied to the new test set without further feature selection or model fitting.

### Feature importance analysis and derivation of a minimal predictor set

To identify chromosomal features contributing to plasmid carriage prediction, impurity-based feature importance scores were extracted from the optimized RF classifier trained on the 2,824-gene signature set. Feature importance was quantified as the mean decrease in Gini impurity^32^ and averaged across five independent training-test splits.

To derive a minimal feature set predictive of plasmid carriage, we implemented an incremental feature selection procedure. Features were first ranked according to their mean Gini importance scores, after which progressively reduced feature sets were generated by retaining only the highest-ranked genes. Random Forest models were retrained on each reduced feature matrix and evaluated using the same five-seed validation framework as the full model. The parsimonious predictor set was defined as the smallest feature subset recovering at least 98% of the predictive performance observed for the full feature set. This procedure identified a minimal set of 15 chromosomal genes.

To assess the generality of this reduced signature, the 15-gene product feature matrix was used to train models for plasmid number prediction, total plasmid size prediction, and plasmid mobility classification. Model training and evaluation followed the same procedures described for the primary analyses.

To examine the phylogenetic distribution of the minimal feature set, the prevalence of top-ranked genes was quantified across prokaryotic genera as the fraction of genomes carrying each gene. Taxonomic similarity among genera was calculated using Gower distance^75^ based on hierarchical taxonomic assignments (superkingdom, phylum, class, order, and family), followed by hierarchical clustering using Ward’s minimum variance method^76^.

### Functional category-specific predictive modeling

To evaluate the relative contributions of different chromosomal functional categories to prediction, the 2,824 chromosomal features were manually grouped according to KEGG BRITE functional categories^77,78^. A curated keyword list was compiled for each BRITE category, and gene-product annotations were assigned to one or more categories by case-insensitive regular-expression matching after removal of non-alphanumeric characters from both product names and keywords. Features annotated as “Unknown” or “Protein of unknown function” were excluded from downstream analyses.

For each functional category, an independent feature matrix was constructed using only genes assigned to that category. RF models were subsequently trained and evaluated using the same five-seed 80/20 train–test framework described above. Performances were compared to identify categories with the greatest predictive power.

Among the evaluated categories, enzyme-associated genes, transporters, polyketide biosynthesis proteins, and non-coding RNAs exhibited the highest predictive performance. Because enzyme-associated genes represented the largest functional module, whereas non-coding RNAs comprised a highly compact feature set, these two categories were subjected to further downstream characterization.

### Enzyme-specific predictive modeling and hierarchical EC analysis

Because enzyme-associated genes exhibited the strongest predictive power among all functional categories, we further investigated the contribution of host enzymatic architecture to plasmid-associated traits. From the 2,824 chromosomal features identified in the primary analysis, 1,121 genes annotated as enzymes were extracted for enzyme-focused modeling. To standardize functional annotations and enable hierarchical analyses, enzyme genes were mapped to Enzyme Commission (EC) identifiers^35^ using RefSeq product annotations. Gene products assigned to the same EC entry were grouped together, and their chromosomal copy numbers were summed to generate genome-level EC abundance profiles. This procedure transformed gene-centric feature matrices into function-centric representations of host metabolic capacity.

The resulting EC profiles were analyzed at multiple levels of the EC hierarchy. Specifically, abundances were aggregated independently at four resolutions corresponding to the four EC levels: specific enzymatic reactions (EC4), sub-subclasses (EC3), subclasses (EC2), and major enzyme classes (EC1). For each genome, separate feature matrices were constructed at each level by summing the abundances of all descendant EC entries belonging to the corresponding parent category.

To evaluate how predictive information is distributed across enzymatic hierarchies, independent RF models were trained using the EC1, EC2, EC3, and EC4 feature matrices following the same training, validation, and hyperparameter optimization procedures described above. Model performance was compared across four resolution levels for plasmid carriage classification and plasmid size regression.

To characterize global variation in host enzymatic architecture, the relative abundances of the seven EC1 enzyme classes—oxidoreductases, transferases, hydrolases, lyases, isomerases, ligases, and translocases—were additionally calculated for each genome. These proportional profiles were used for comparative analyses across taxonomic groups and for proportion-based predictive modeling.

Feature importance was quantified as the mean decrease in Gini impurity for RF classifiers and as the mean decrease in squared-error impurity (MDI)^32^ for RF regressors. For the plasmid-size regression models, EC4-level MDI scores were aggregated according to their parent EC3 categories and compared with the corresponding MDI scores obtained directly from EC3-based regressors. Concordance between the two estimates was quantified using Pearson correlation analysis.

### Prediction of plasmid functional cargo and antibiotic resistance genes

To investigate whether host chromosomal architecture encodes information about plasmid-borne functional repertoires, we first quantified the abundance of major functional gene categories encoded on plasmids. Product annotations from plasmid replicons were screened against curated reference sets representing four functionally important groups: toxins, transposases, transcription factors, and prokaryotic defense systems. For each genome, the abundance of plasmid-encoded genes belonging to each category was calculated and used as quantitative output variables in downstream analyses.

Plasmid-encoded antibiotic resistance genes (ARGs) were annotated using the Resistance Gene Identifier (RGI, version 6.0.5) and the Comprehensive Antibiotic Resistance Database (CARD, version 4.0.1)^36^. Only annotations meeting the CARD “Strict” criteria were retained. Identified ARGs were subsequently classified into 33 resistance categories according to their resistance mechanisms. For each genome, the abundance of genes belonging to each ARG category was quantified separately.

To evaluate whether chromosomal gene content predicts plasmid functional composition, RF regression models were trained using multiple chromosomal feature representations, including the 2,824-gene chromosomal signature, the 1,121-gene enzyme subset, the 15-gene minimal predictor set, the 12-feature non-coding RNA panel, and EC-based enzyme representations. For ARG-specific analyses, models were further trained for each resistance category using either the 2,824-gene chromosomal feature set or the EC4 reaction-level enzyme representation. Analyses were restricted to plasmid-containing genomes. Model performance was evaluated using five-fold cross-validation with shuffled. To assess whether predictive relationships between chromosomes and plasmid functional cargo were maintained within individual bacterial lineages, model performance was also evaluated within the genera *Escherichia* and *Salmonella*, the two most abundant genera in the dataset.

## Data Availability

All the data associated with this work are available at the GitHub repository (https://github.com/hyyy0187/plasmid-biology-prediction).

## Code Availability

All codes are available at the GitHub repository (https://github.com/hyyy0187/plasmid-biology-prediction).

## Acknowledgement

This study was supported by the National Key R&D Program of China (2024YFA0920200 to TW), the National Natural Science Foundation of China (12401660 and 32470701 to TW), and the Shenzhen Institute of Synthetic Biology Scientific Research Program (HSE499011086 to TW). We are grateful to the Shenzhen Infrastructure for Synthetic Biology for providing instrument support and technical assistance.

