## Supplementary Information for "Plasmid biology is compressed in host chromosomal architecture"

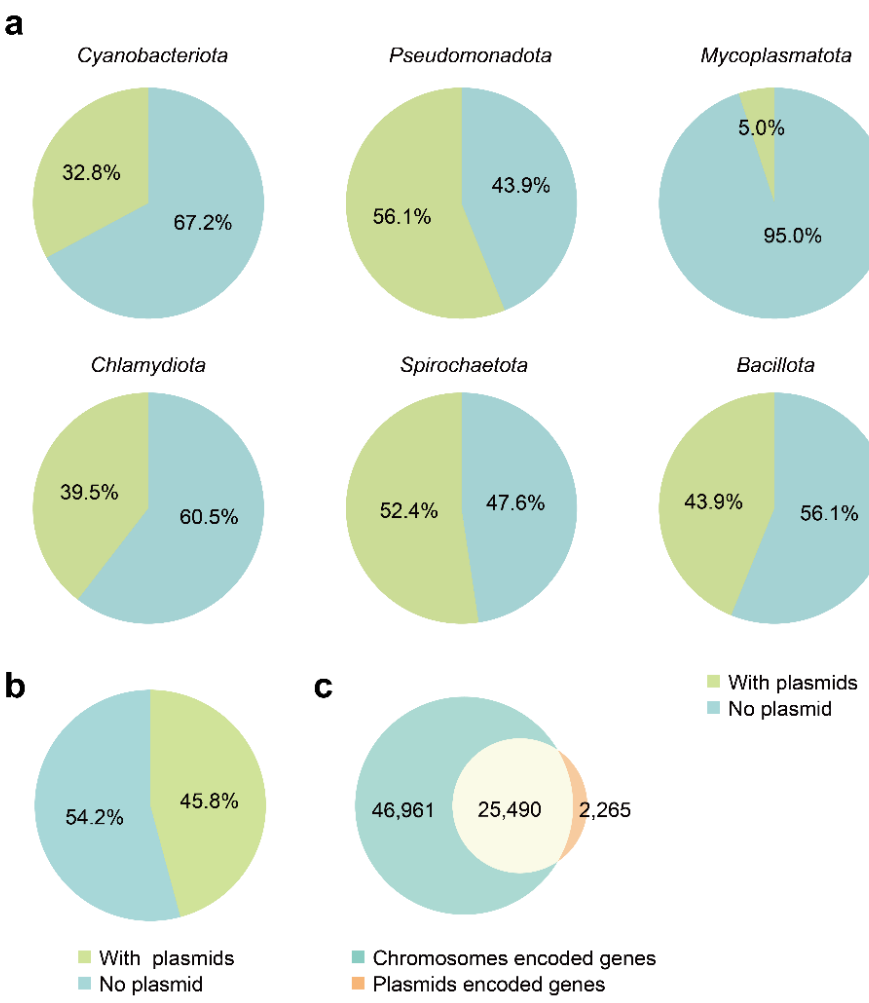


**Fig. S1| Plasmid distribution and gene repertoire across prokaryotic genomes**

(a) Pie charts show the proportion of plasmid-carrying (green) and plasmid-free (blue) genomes across six representative phyla.

(b) The fractions of prokaryotic genomes with and without plasmids.

(c) Venn diagram showing the functional overlap between chromosomal and plasmid-encoded gene repertoires.

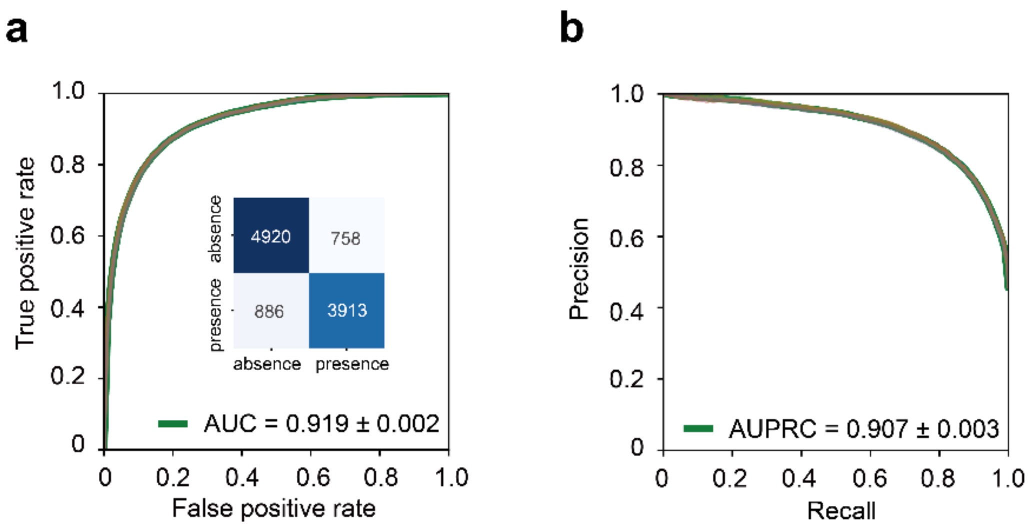


**Fig. S2| Performance of the random forest model for plasmid carriage prediction using 2,824 chromosomal gene features.**

(a) Prediction of plasmid carriage from chromosomal gene features. Performance metrics were obtained from five independent random forest model refits using stratified 80/20 train–test splits generated with different random seeds and are reported as mean ± s.d. The ROC curve shows the mean true positive rate across the five replicates, with individual replicate shown as faint lines and the shaded region indicating ± s.d.; the mean AUROC ± s.d. across all five refits is reported in the legend. Inset, element-wise mean confusion matrix across the five refits, with values truncated to integers.

(b) Precision–recall curves for the same model across the five independent random forest model refits. The precision–recall curve shows the mean precision across the five replicates, with individual replicate shown as faint lines and the shaded region indicating ± s.d.; the mean AUPRC ± s.d. across all five replicates is reported in the legend.

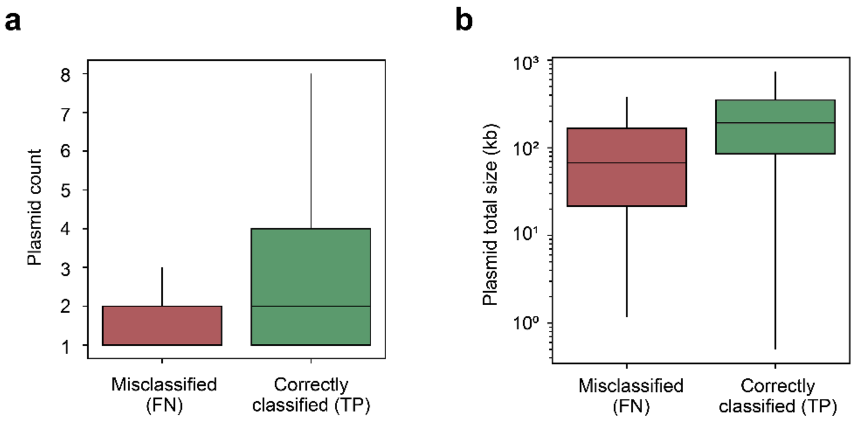


**Fig. S3| Misclassified genomes exhibit marginal plasmid content.**

(a) Boxplots compare plasmid number per genome between misclassified (false negatives, n = 4,402 pooled across five test splits) and correctly classified (true positives, n = 19,598 pooled) plasmid-carrying genomes. False negatives contain significantly fewer plasmids than true positives. Boxplots show median (center line), interquartile range (box limits), and whiskers extending to minimum and maximum values excluding outliers. Statistical significance was assessed using a one-sided Mann–Whitney U test ($P < 0.001$).

(b) Boxplots compare plasmid total size per genome between misclassified and correctly classified plasmid-carrying genomes. Boxplot definitions and statistical analyses are as described in a.

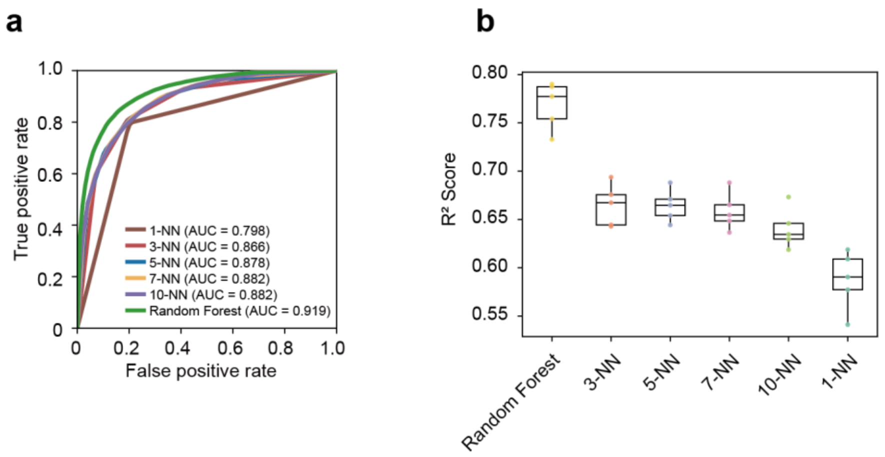


**Fig. S4| Comparison of Random Forest and distance-based *k*-nearest neighbor (k-NN) models.**

(a) ROC curves for plasmid carriage classification. Random Forest achieved the highest AUROC, outperforming all k-NN models (k = 1, 3, 5, 7, 10).

(b) Prediction of total plasmid size. Boxplots show test-set $R^2$ values across five independent train–test splits. Center lines indicate medians, boxes denote interquartile ranges, and whiskers indicate minimum and maximum values; individual replicate scores are overlaid as dots.

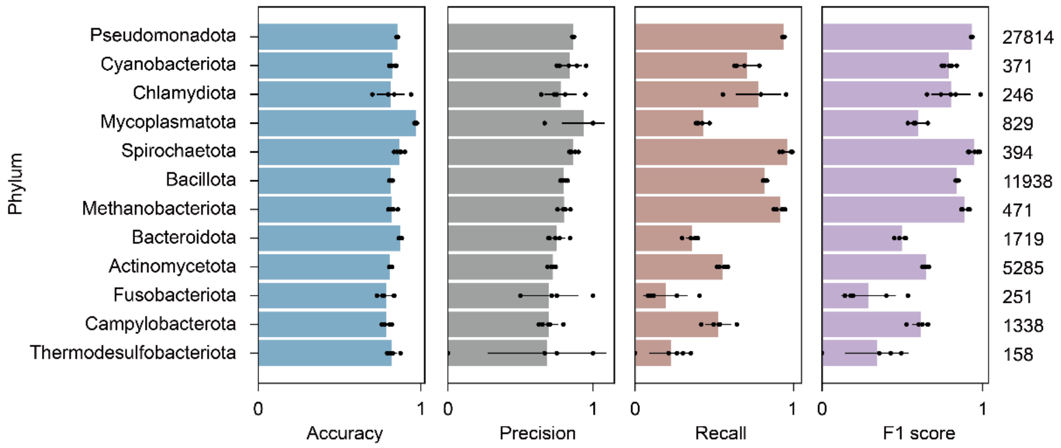


**Fig. S5| Phylum-level performance of the plasmid carriage classifier.** Horizontal bar plots show mean accuracy, precision, recall, and F1 score across five stratified train–test splits for phyla represented by at least 100 test genomes. Bars and error bars denote mean ± s.d.; numbers indicate cumulative number of test-set genomes across five splits.

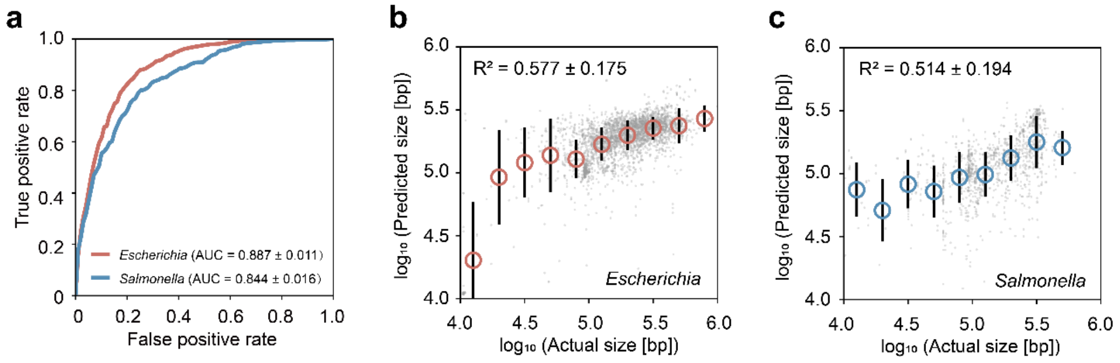


**Fig. S6| Genus-specific performance of the prediction models in *Escherichia* and *Salmonella*.**

(a) ROC curves for plasmid carriage prediction within *Escherichia* and *Salmonella* test subsets. Each curve represents the mean performance across five independent train–test splits.

(b, c) Genus-specific prediction of total plasmid size in *Escherichia* (b) and *Salmonella* (c). Scatter plots show observed versus predicted total plasmid size ($\log_{10}$ bp). Grey points represent individual genomes. Circles (*Escherichia*, red; *Salmonella*, blue) indicate mean predicted values within ten equal-width bins of observed plasmid size, and error bars denote ± s.d. within each bin. The $R^2$ value shown in each panel represents the mean predictive performance across five independent train–test splits.

resents the mean predictive performance across five independent train–test splits.

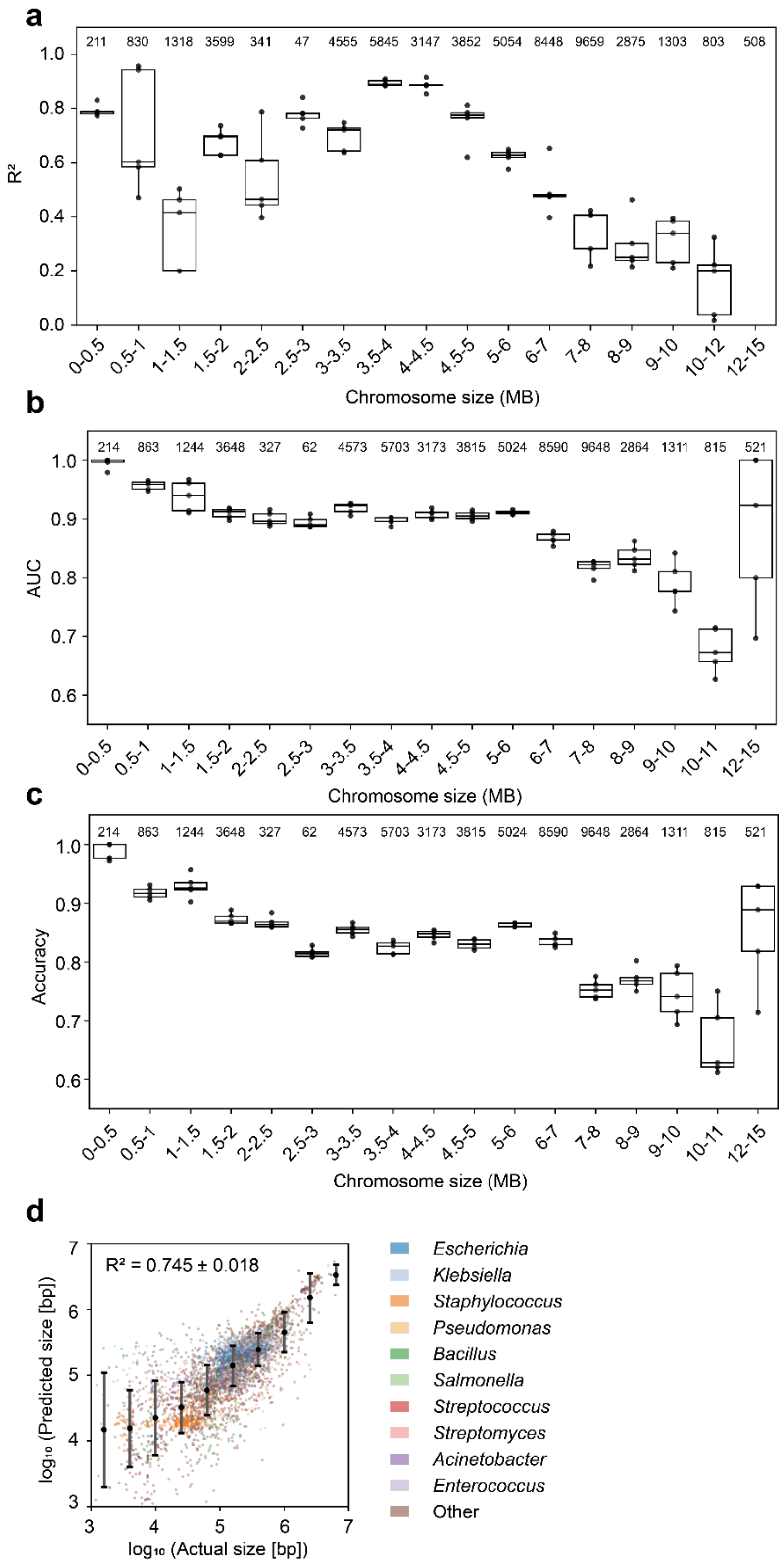


**Fig. S7| Performance and robustness evaluation of the predictive models across varying host chromosome sizes.** Test-set genomes were stratified into 18 host chromosome size bins spanning less than 0.5 to greater than 15 megabases, and model performance was evaluated separately within each bin across five independent replicates.

(a) Per-bin regression performance ($R^2$) for prediction of total plasmid size.

(b, c) Per-bin classification performance for plasmid carriage prediction, measured by AUC (b) and accuracy (c). In all panels, boxplots show the median (center line), interquartile range (box limits), and min–max values (whiskers) across five replicates, with individual replicate values overlaid as dots. Numbers above each bin indicate the cumulative number of test-set genomes evaluated across all five splits.

(d) Random Forest prediction of total plasmid size using chromosome-size-normalized feature gene abundances. Observed versus predicted total plasmid sizes ($\log_{10}$ bp) are shown for genomes in a representative test partition (n = 10,479). Points represent individual genomes and are colored by the ten most abundant bacterial genera (all remaining genera are grouped as "Other"). Filled circles indicate the mean predicted value within bins of observed plasmid size, and error bars denote ± s.d. of predicted values within each bin. The reported $R^2$ represents the mean ± s.d. across five independent train–test splits.

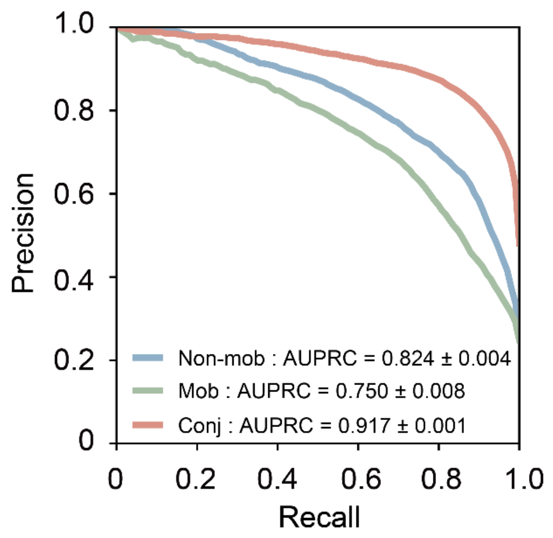


**Fig. S8| Precision–recall performance of the mobility prediction model.** A random forest classifier was trained using 2,824 chromosomal gene features to predict plasmid mobility class, including non-mobilizable, mobilizable, and conjugative plasmids. Precision–recall curves show the mean performance across five stratified train–test splits using different random seeds.

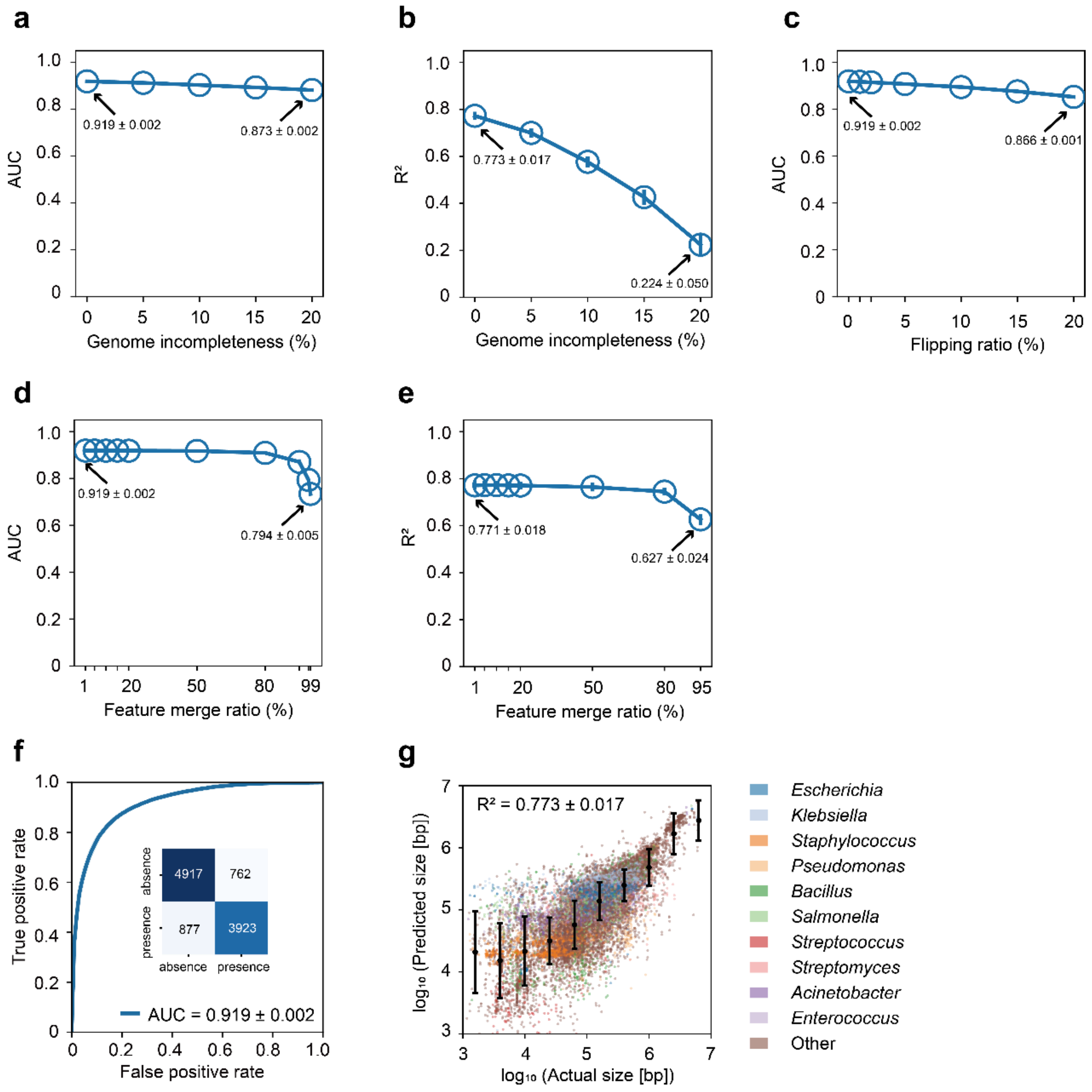


**Fig. S9| Robustness of the predictive models against data perturbations and alternative feature representations.** All metrics are evaluated across five independent train-test splits.

(a, b) Effects of simulated genome incompleteness on plasmid carriage classification (a; AUC) and total plasmid size prediction (b; $R^2$). Gene records were randomly removed from chromosomal feature profiles at rates of 0–20%. Points and error bars represent mean ± s.d. across five replicates

(c) Classification robustness to label flipping ratio. Mean AUC is shown as a function of random training-label flipping (0–20%). Points and error bars represent mean ± s.d. across five replicates

(d, e) Effects of feature redundancy and gene nomenclature inconsistency. Feature columns were randomly merged at increasing proportions to simulate nomenclature collapse. Mean AUC (d) and $R^2$ (e) are shown across merge ratios. Points and error bars represent mean ± s.d. across five replicates.

(f) Performance of a classifier trained on Pfam-domain representations. Gene-product features were mapped to UniProt Pfam domains prior to model training. ROC curves show the mean performance across five replicates.

(g) Prediction of total plasmid size using Pfam-domain features. Scatter plots show observed versus predicted plasmid sizes ($log_{10}$ bp) for a representative test partition. Points represent individual genomes and are colored by the ten most abundant genera. Filled circles indicate mean predicted values within bins of observed plasmid size, and error bars denote ± s.d. of predicted values within each bin. The $R^2$ value shown in the panel represents the mean ± s.d. across five independent train–test splits.

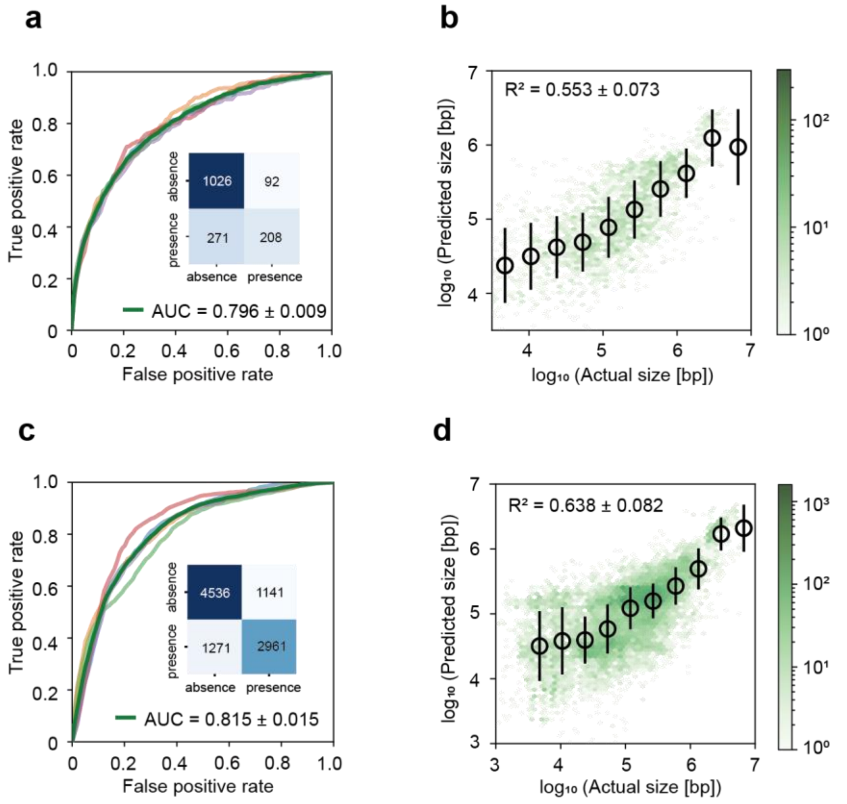


**Fig. S10| Generalizability of the prediction models across taxonomic boundaries.**

(a) ROC curves for plasmid carriage prediction after species-level dereplication, where a single representative genome was retained per species before train–test partitioning. Curves represent the mean performance across five replicates; the inset shows the element-wise mean confusion matrix across the five corresponding train–test splits, with values truncated to integers.

(b) Regression performance for total plasmid size following species-level dereplication. Hexbin plots show observed versus predicted plasmid sizes ($\log_{10}$ bp), with color intensity indicating point density on a $\log_{10}$ scale. Black circles indicate mean predicted values within ten equal-width bins of observed size, and error bars denote ± s.d. The overall $R^2$ is shown in the panel.

(c) ROC curves for plasmid carriage prediction under a strict species-holdout strategy, in which 20% of species were excluded from training and used exclusively for testing. Curves represent the mean performance across five replicates; the inset shows the element-wise mean confusion matrix across the five corresponding train–test splits, with values truncated to integers.

(d) Regression performance for total plasmid size under the species-holdout strategy. Hexbin plots show observed versus predicted plasmid sizes ($\log_{10}$ bp), with color intensity representing point density. Black circles and error bars denote the binned mean ± s.d. across ten equal-width bins.

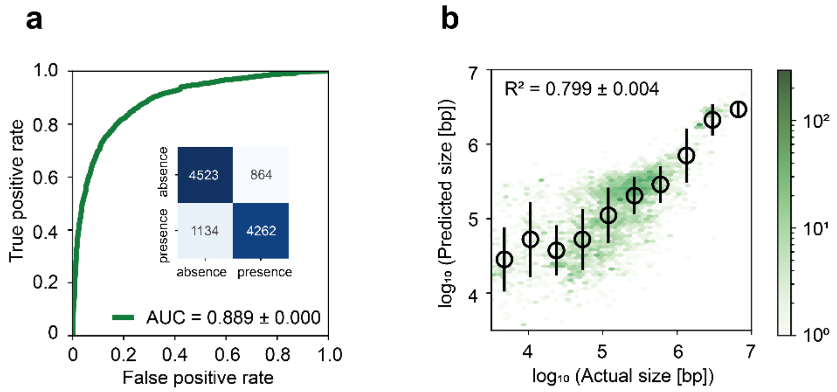


**Fig. S11| Validation on an independent genome collection**

(a) Prediction of plasmid carriage from chromosomal gene-product composition on an independent external test set comprising 10,784 prokaryotic genomes. Performance metrics were obtained from five independent random forest model refits and are reported as mean ± s.d. The ROC curve shown is from a representative refit (seed 42), with the mean AUC ± s.d. across all five refits reported in the legend. Inset, element-wise mean confusion matrix across the five refits, with values truncated to integers.

(b) Prediction of total plasmid size from chromosomal gene-product composition on the same external test set. Density scatterplot (hexagonal binning) of predicted versus observed plasmid size ($\log_{10}$ bp) for a representative model refit; color scale, $\log_{10}$(point density). Mean $R^2$ ± s.d. across the five model refits is reported in the panel.

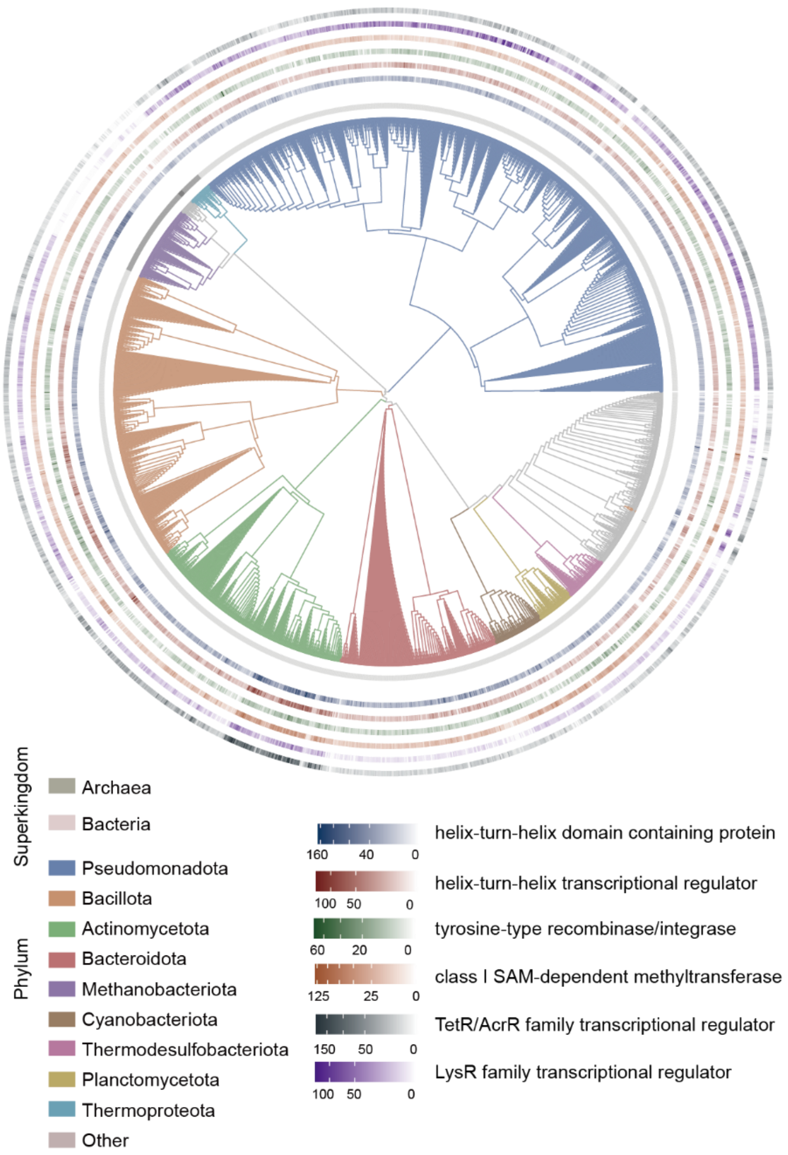


**Fig. S12| Taxonomic distribution of the six highest-ranking chromosomal features associated with plasmid carriage.** The taxonomy-based circular dendrogram illustrates the genus-level distribution of the six chromosomal features with the highest mean random-forest feature importance for plasmid carriage classification. Feature abundance was quantified as the mean per-genome copy number within each genus represented in the dataset. The dendrogram was generated by hierarchical clustering using Ward D2 linkage on pairwise Gower distances calculated from five taxonomic levels: superkingdom, phylum, class, order and family. Genus names were used only as leaf labels and were not included in the distance calculation. Each tip represents a single genus, with branch and tip colors indicating the nine phyla containing the largest numbers of genera; all remaining phyla are grouped as "Other" (grey). From the inside out, the concentric tracks show superkingdom assignment (Bacteria or Archaea), followed by six heatmap tracks representing the six chromosomal features. Color intensities were scaled independently for each feature according to its mean copy number, highlighting taxon-associated variation in the abundance of these high-ranking features.

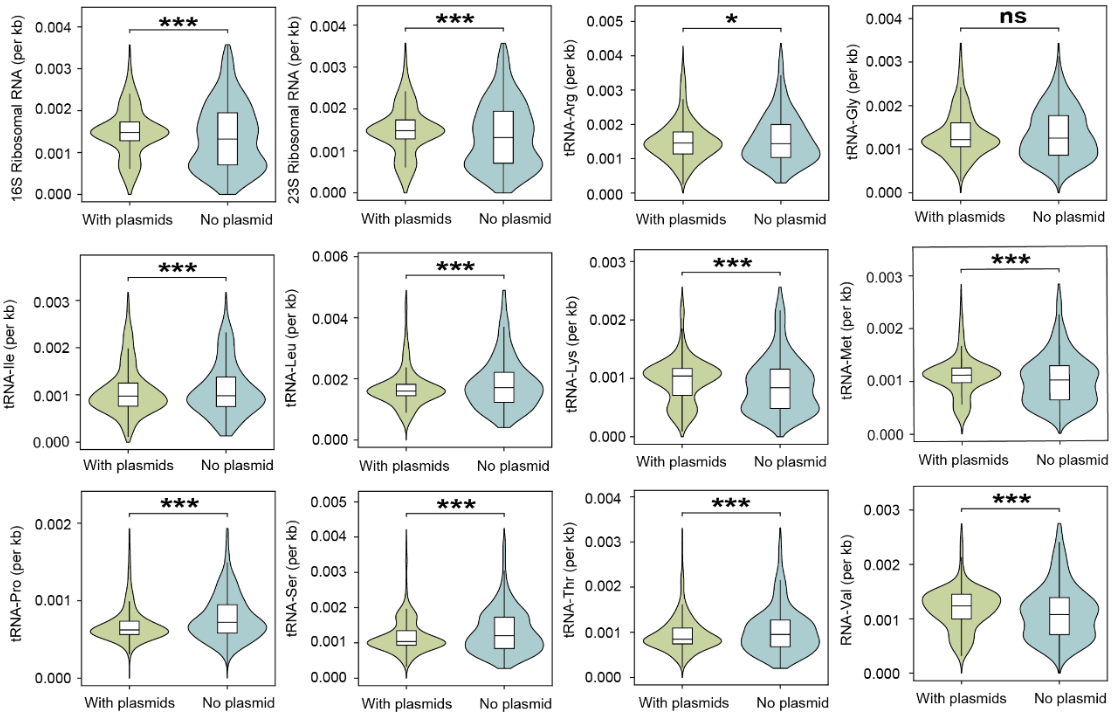


**Fig. S13| Distributions of 12 ncRNA-associated chromosomal features in genomes with versus without plasmids.** Each panel shows one ncRNA feature normalized by chromosome length (counts per kb), stratified by plasmid carriage status ("No plasmid" vs "With plasmids"). Split violins (kernel density, bandwidth-adjusted) show per-kb distributions, with overlaid boxplots indicating median and interquartile range (whiskers extend to 1.5× IQR; outliers not shown). Values above the 99th percentile were truncated for visualization, while statistical tests were performed on the full untrimmed data. Group differences were assessed using two-sided Mann–Whitney U tests. Asterisks indicate significance level (*$P < 0.05$, **$P < 0.01$, ***$P < 0.001$; ns, not significant).

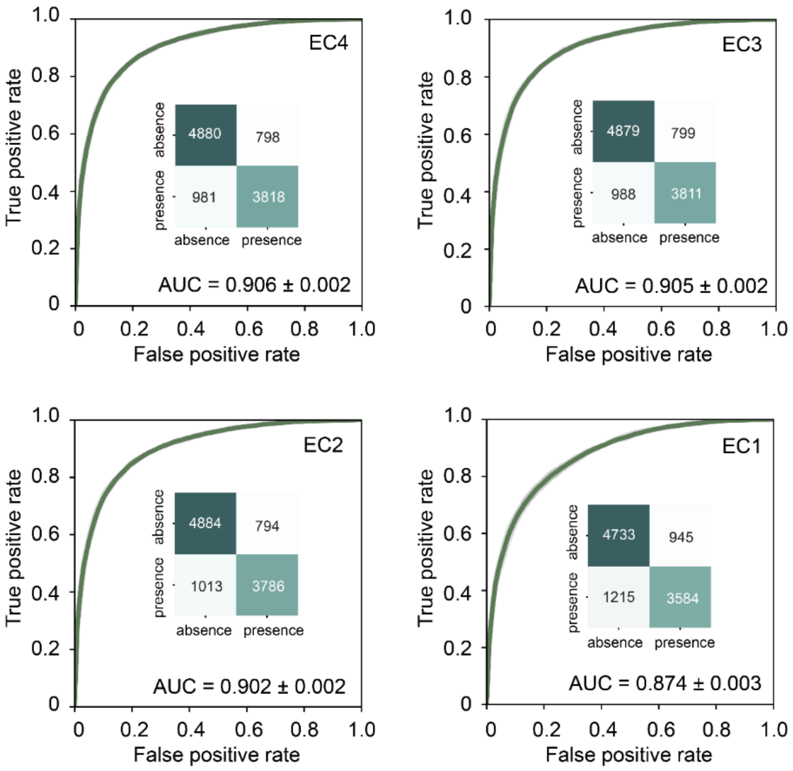


**Fig. S14| Performance of plasmid carriage classification across Enzyme Commission (EC) hierarchy levels.** Random Forest models were trained using chromosomal enzymatic features aggregated from EC4 to EC1. ROC curves represent mean performance across five independent train–test splits. The inset shows the element-wise mean confusion matrix across the five splits, with values rounded to integers.

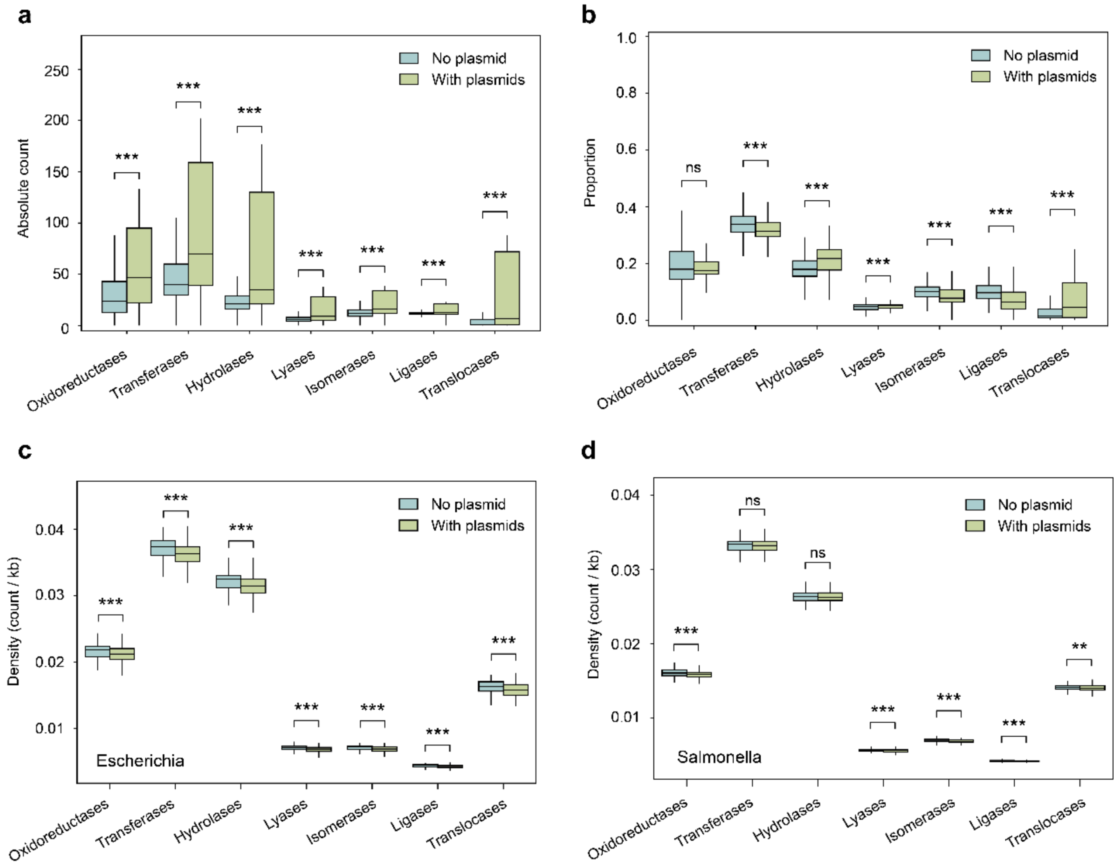


**Fig. S15| Chromosomal enzyme composition associated with plasmid carriage at global and genus-specific levels.**

(a) Absolute chromosomal abundances of the seven EC1 classes in plasmid-free and plasmid-bearing genomes. Enzyme abundance was defined as the number of chromosome-encoded enzymes assigned to each EC1 class. Boxes indicate the median and interquartile range, and whiskers extend to 1.5 times the interquartile range; outliers are not shown. Differences between plasmid-free and plasmid-bearing genomes were assessed using two-sided Mann–Whitney U tests, with Benjamini–Hochberg false-discovery-rate correction across the seven enzyme classes. (***, $q < 0.001$; **, $q < 0.01$; *, $q < 0.05$; ns, not significant.)

(b) Relative chromosomal composition of the seven EC1 classes. For each genome, the abundance of each EC1 class was expressed as a proportion of the total number of EC-classified chromosomal enzymes. Boxplot definitions and statistical analyses are as described in a.

(c) Chromosomal enzyme densities in plasmid-free (n = 644) and plasmid-bearing (n = 3,559) *Escherichia* genomes. Enzyme density was calculated as the number of chromosome-encoded enzymes assigned to each EC1 class per kilobase of chromosomal sequence. Statistical analyses are as described in a.

(d) Chromosomal enzyme densities in plasmid-free (n = 554) and plasmid-bearing (n = 1,299) *Salmonella* genomes. Enzyme densities and statistical analyses are as described in c.

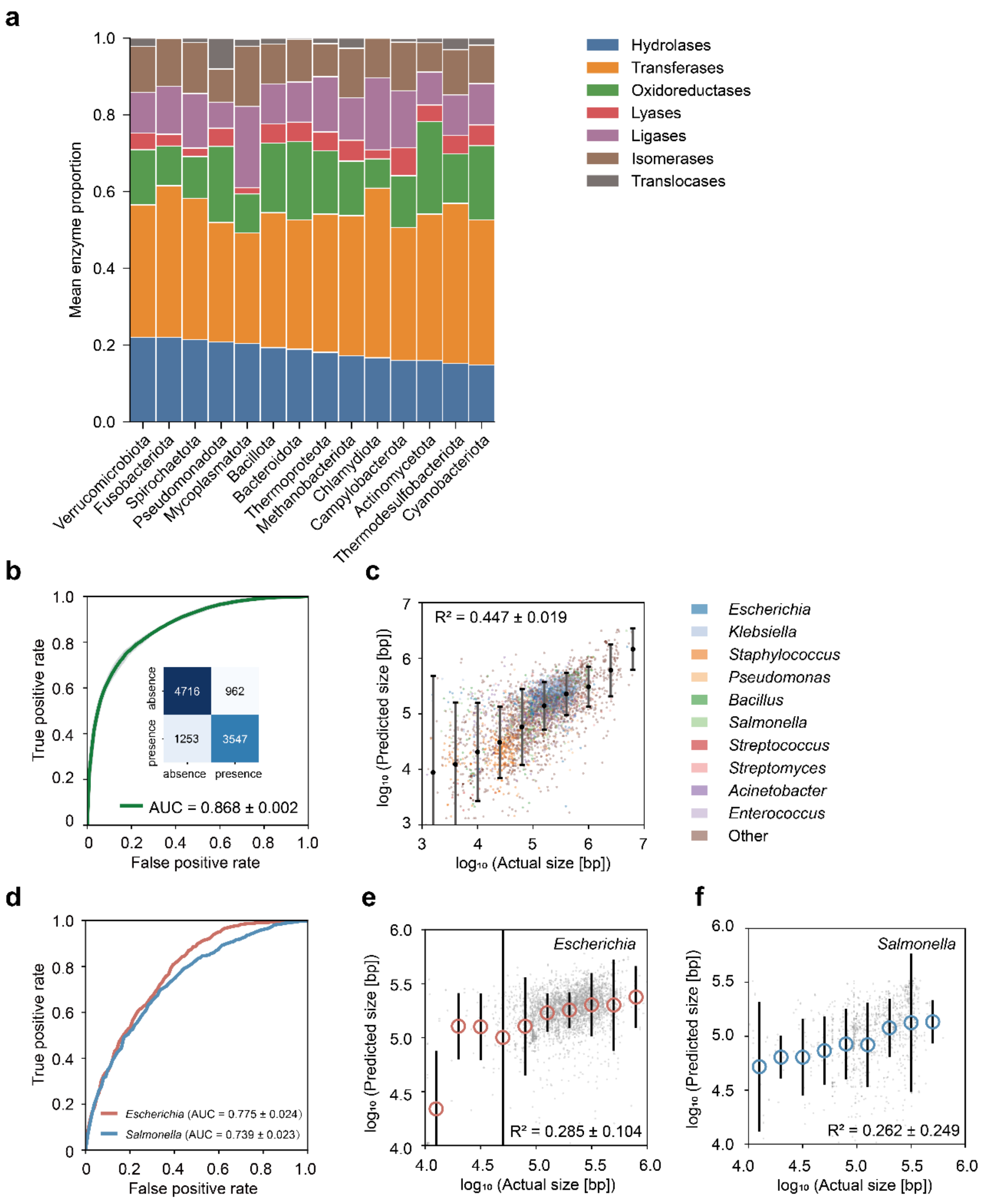


**Fig. S16| Chromosomal enzyme-class composition and its predictive capacity for plasmid carriage and total plasmid size.**

(a) Phylum-level EC1 composition of chromosomal enzymes. For each genome, EC1 proportions were computed and averaged within phyla (≥100 genomes per phylum).

(b) Prediction of plasmid carriage from EC1 composition. Random Forest classifier performance was evaluated across five independent train–test splits. ROC curves show individual replicates (thin grey lines), mean performance (green line), and ± s.d. shading; mean AUC ± s.d. is reported in the legend. The inset shows the element-wise mean confusion matrix across splits, with values rounded to integers.

(c) Prediction of total plasmid size using EC1 composition. Points represent individual genomes, colored by the ten most abundant genera (others grouped as "Other"). Models use chromosomal EC1 proportions across seven enzyme classes (oxidoreductases, transferases, hydrolases, lyases, isomerases, ligases, and translocases). Filled circles indicate binned mean predictions, with error bars showing ± s.d. within bins. $R^2$ values are reported as mean ± s.d. across five independent train–test splits.

(d) Genus-specific prediction of plasmid carriage from EC1 composition. Mean ROC curves are shown for *Escherichia* and *Salmonella* across five independent train–test splits. AUROC values are reported as mean ± s.d. across splits.

(e) Prediction of total plasmid size in *Escherichia* using EC1 composition. Points show pooled test-set predictions across five independent train–test splits. Filled circles indicate binned mean predictions, with error bars showing ± s.d. within bins. $R^2$ is reported as mean ± s.d. across splits.

(f) Prediction of total plasmid size in *Salmonella* using EC1 composition. Points show pooled test-set predictions across five independent train–test splits. Filled circles indicate binned mean predictions, with error bars showing ± s.d. within bins. $R^2$ is reported as mean ± s.d. across splits.

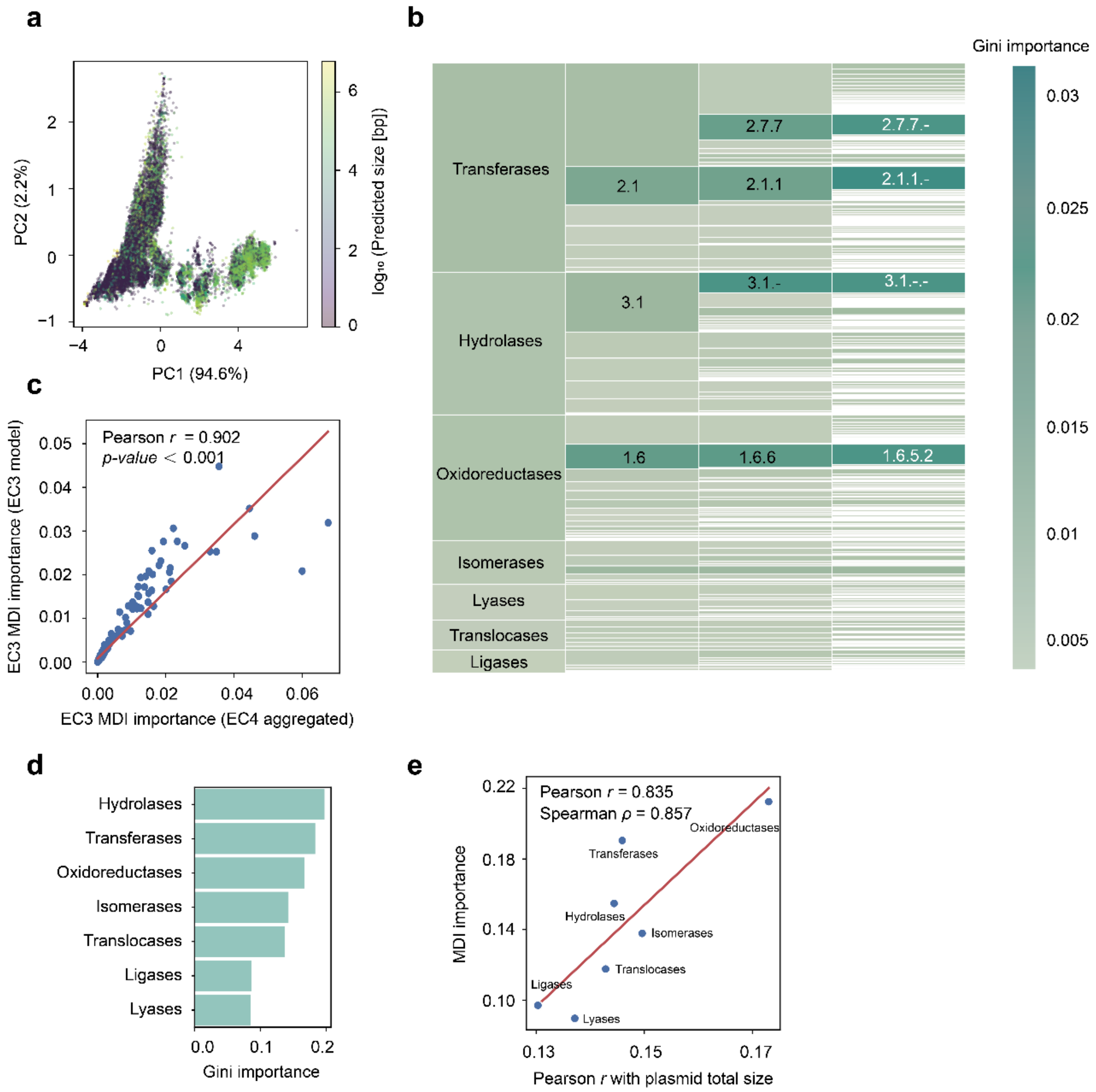


**Fig. S17| Multivariate organization and feature importance of chromosomal enzyme architecture across EC levels.**

(a) Principal component analysis of chromosomal EC1 profiles in relation to total plasmid size. PCA was performed on the standardized chromosomal abundances of the seven EC1 enzyme classes. Each point represents one genome and is colored according to its observed $\log_{10}$-transformed total plasmid size. Axis labels indicate the proportion of total variance explained by the first two principal components.

(b) Hierarchical distribution of Gini importance across EC levels for plasmid-carriage classification. EC4-level Gini importance scores were averaged across five runs. At the EC4 level, both rectangle area and color intensity represent mean Gini importance. At higher levels (EC3–EC1), rectangle area represents cumulative Gini importance aggregated from descendant EC4 features, whereas color intensity represents the Gini-importance-weighted mean of the corresponding descendant EC4 scores.

(c) Concordance of mean decrease in impurity (MDI) scores between EC3- and EC4-resolution Random Forest regressors for plasmid-size prediction. EC4-level MDI scores were summed within EC3 categories and compared with mean MDI scores obtained directly from EC3-based regressors. Each point represents an EC3 category. The red line shows the linear fit. Pearson's $r$ and $p$-value are reported.

(d) Feature importance of the seven EC1 classes for predicting plasmid carriage. Importance scores were averaged across five train–test splits, with bars arranged in descending order.

(e) Relationship between Random Forest feature importance and marginal feature–target association. Each point represents one EC1 enzyme class. The x axis shows Pearson's correlation coefficient between the abundance of each EC1 feature and untransformed total plasmid size, whereas the y axis shows its mean decrease in impurity (MDI) importance averaged across five independent Random Forest regressors. The red line represents an ordinary least-squares fit. Pearson's $r$ and Spearman's $\rho$ quantify the linear and rank-based concordance, respectively, between feature–target correlation and model-derived importance across the seven EC1 classes.

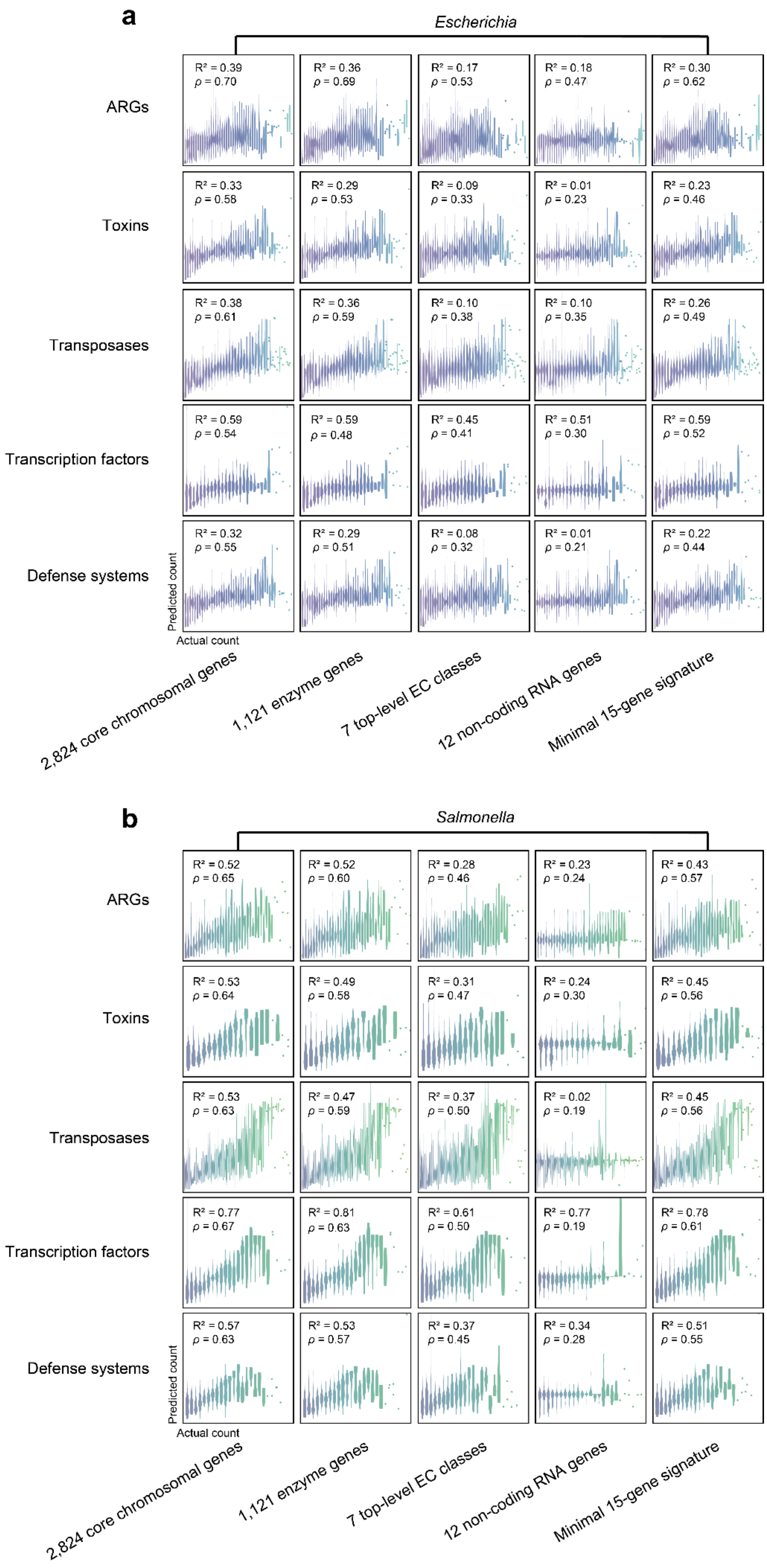
a
Escherichia
ARGs
Toxins
Transposases
Transcription factors
Defense systems
$R^2$ = 0.39 $\rho$ = 0.70
$R^2$ = 0.36 $\rho$ = 0.69
$R^2$ = 0.17 $\rho$ = 0.53
$R^2$ = 0.18 $\rho$ = 0.47
$R^2$ = 0.30 $\rho$ = 0.62
$R^2$ = 0.33 $\rho$ = 0.58
$R^2$ = 0.29 $\rho$ = 0.53
$R^2$ = 0.09 $\rho$ = 0.33
$R^2$ = 0.01 $\rho$ = 0.23
$R^2$ = 0.23 $\rho$ = 0.46
$R^2$ = 0.38 $\rho$ = 0.61
$R^2$ = 0.36 $\rho$ = 0.59
$R^2$ = 0.10 $\rho$ = 0.38
$R^2$ = 0.10 $\rho$ = 0.35
$R^2$ = 0.26 $\rho$ = 0.49
$R^2$ = 0.59 $\rho$ = 0.54
$R^2$ = 0.59 $\rho$ = 0.48
$R^2$ = 0.45 $\rho$ = 0.41
$R^2$ = 0.51 $\rho$ = 0.30
$R^2$ = 0.59 $\rho$ = 0.52
$R^2$ = 0.32 $\rho$ = 0.55
$R^2$ = 0.29 $\rho$ = 0.51
$R^2$ = 0.08 $\rho$ = 0.32
$R^2$ = 0.01 $\rho$ = 0.21
$R^2$ = 0.22 $\rho$ = 0.44
Predicted count
Actual count
2,824 core chromosomal genes
1,121 enzyme genes
7 top-level EC classes
12 non-coding RNA genes
Minimal 15-gene signature
b
Salmonella
ARGs
Toxins
Transposases
Transcription factors
Defense systems
$R^2$ = 0.52 $\rho$ = 0.65
$R^2$ = 0.52 $\rho$ = 0.60
$R^2$ = 0.28 $\rho$ = 0.46
$R^2$ = 0.23 $\rho$ = 0.24
$R^2$ = 0.43 $\rho$ = 0.57
$R^2$ = 0.53 $\rho$ = 0.64
$R^2$ = 0.49 $\rho$ = 0.58
$R^2$ = 0.31 $\rho$ = 0.47
$R^2$ = 0.24 $\rho$ = 0.30
$R^2$ = 0.45 $\rho$ = 0.56
$R^2$ = 0.53 $\rho$ = 0.63
$R^2$ = 0.47 $\rho$ = 0.59
$R^2$ = 0.37 $\rho$ = 0.50
$R^2$ = 0.02 $\rho$ = 0.19
$R^2$ = 0.45 $\rho$ = 0.56
$R^2$ = 0.77 $\rho$ = 0.67
$R^2$ = 0.81 $\rho$ = 0.63
$R^2$ = 0.61 $\rho$ = 0.50
$R^2$ = 0.77 $\rho$ = 0.19
$R^2$ = 0.78 $\rho$ = 0.61
$R^2$ = 0.57 $\rho$ = 0.63
$R^2$ = 0.53 $\rho$ = 0.57
$R^2$ = 0.37 $\rho$ = 0.45
$R^2$ = 0.34 $\rho$ = 0.28
$R^2$ = 0.51 $\rho$ = 0.55
Predicted count
Actual count
2,824 core chromosomal genes
1,121 enzyme genes
7 top-level EC classes
12 non-coding RNA genes
Minimal 15-gene signature

**Fig. S18| Genus-resolved prediction of plasmid functional cargo using different chromosomal feature sets.** Prediction performance is shown separately for *Escherichia* (a) and *Salmonella* (b). Columns correspond to five chromosomal feature sets (2,824 core chromosomal genes, 1,121 enzyme genes, seven EC1 classes, 12 ncRNA features, and a minimal 15-gene signature), and rows represent five plasmid functional cargo classes (antibiotic resistance genes, toxins, transposases, transcription factors, and prokaryotic defense systems). Within each panel, observed values are plotted against predicted values. The corresponding $R^2$ and Spearman's $\rho$ are shown in each panel. Predicted values are out-of-fold (OOF) predictions from 5-fold cross-validation; the reported $R^2$ was calculated by pooling out-of-fold predictions across all folds.
